# Integrating Economic dynamics into Ecological Networks: The case of fishery sustainability

**DOI:** 10.1101/840314

**Authors:** Paul Glaum, Valentin Cocco, Fernanda S. Valdovinos

## Abstract

Understanding and sustainably managing anthropogenic impact on ecosystems requires studying the integrated economic -ecological dynamics driving coupled human-natural systems. Here, we expand ecological network theory to study fishery sustainability by incorporating economic drivers into food-web models to evaluate the dynamics of thousands of single-species fisheries across hundreds of generated food-webs and two management strategies. Analysis reveals harvesting high population biomass species can initially support fishery persistence, but threatens long term economic and ecological sustainability by indirectly inducing extinction cascades in non-harvested species. This dynamic is exacerbated in open access fisheries where profit driven growth in fishing effort increases perturbation strength. Results demonstrate the unique insight into both ecological dynamics and sustainability garnered from considering economically dynamic fishing effort in the network.

**One Sentence Summary:** Integrating economic drivers into ecological networks reveal non-linear drivers of sustainability in fisheries.

## Main

The advent of network theory in ecology and environmental studies has greatly advanced the study of ecological dynamics and complexity (1, 2). These advances have also translated into a growing knowledge of how human caused disturbances can create far-reaching ecological impacts through indirect effects (3-6). Often, however, the disturbances in these network studies have been studied as a one-time event or a constant external rate of change, separate from the dynamic elements of the network (3, 6, 7). We argue that developing both sustainable management practices and a fuller understanding of anthropocene ecological dynamics require recognition that much of the ecological impact of anthropogenic activity is determined by an integrated feedback process between ecological dynamics and socio-economic conditions as a coupled natural-human system (8, 9). In short, the scope of ecological networks should contain humans as dynamic elements when necessary (10, 11). Here, we expand ecological network theory by incorporating economic dynamics into food-web models to evaluate the coupled natural-human dynamics affecting sustainability in the case of fisheries.

Fisheries are an important example of natural-human integrated network systems given their critical role in the economic stability and food security of billions of people (12). Both the available yield and mobilized fishing effort in any fishery are products of a complex set of interacting socio-economic and ecological factors (13, 14). Understanding the dynamics and consequences of these interactions is particularly pressing as current management strategies have produced an over-exploitation crisis in a multitude of fisheries across the globe (15, 16), threatening both aquatic biodiversity (17) and the aforementioned food security (12). Traditionally, fishery sustainability goals have been implemented into policy on a single-fishery basis as a yield optimization process called maximum sustainable yield (MSY) (12). However, MSY’s conventional consideration of harvested species in isolation, instead of part of a broader food-web (14, 18, 19), has limited its ability to address the indirect effects of harvesting seen in more network based approaches. Specifically, the population variability induced directly through fishing effort on harvested species (20) can transfer to non-harvested species through trophic interactions (5, 21). This variability has been linked to perturbations causing reductions in aquatic biodiversity (22, 23) and ecosystem function (24). These reductions cycle back to further affect harvested species (5, 25) and the potential economic returns harvested species provide (15, 26). Any effect on economic returns can impact future fishing effort and the corresponding effect on harvested ecosystems, creating a bio-economic feedback loop. While this bio-economic loop is understood in concept, limited work exists on its actual dynamics, especially in the context of broader ecological networks (27). We propose that attempts to understand and manage fishery dynamics must consider economic factors in the network as integrated ecological economic models (13). Otherwise, until more is known about the complex interactions between ecological and economic factors driving fishing efforts, policy makers risk attempting to optimize a process we poorly understand.

In accordance with such concerns, we applied network theory to incorporate the ecological complexity of species trophic interactions into the model. This differs from past approaches using Ecopath with Ecosim (28) which rely on extensive lists of system- and species-specific parameters to create models to effectively manage specific systems. The network approach allows for the development of more fundamental, widely-applicable theory because network models can be run across an array of ecological configurations, both empirical (5) and realistically generated (29), by parameterizing metabolism and species interactions through allometry (30). In this study we generated networks as food webs using the Niche Model (31), each with an initial 30 interacting trophic species. Species available for harvest are labeled “fish” for ease of description (Fig 1a). Ecological dynamics in each web were governed by a series of ordinary differential equations and parameterized through allometrically scaled rates (30) creating Allometric-Trophic Network (ATN) models (see Methods, 32). Finally, the network approach also provides a flexible framework which facilitates the integration of economic dynamics.

**Figure 1:**
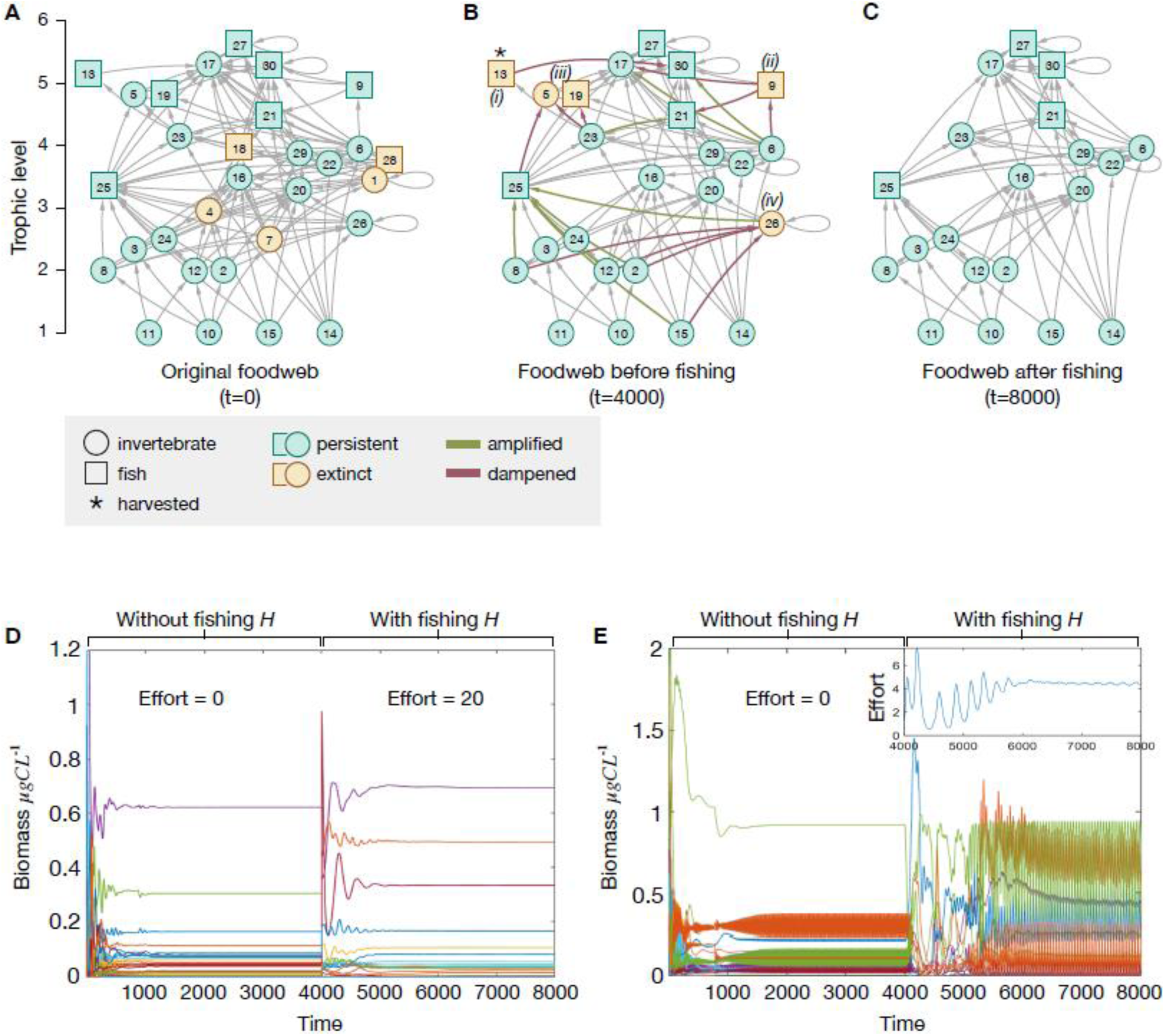
Diagrams of the experimental design. a-c) Example evolution of a food web structure across complete simulation process with a Fixed Effort of 5. Node sizes are logarithmically scaled to the biomass at each point in time of the simulation. Edges between nodes represent trophic interactions with arrows indicating the consumer. The vertical axis indicates the trophic level of trophic species. a) The example food web at initialization (t=0). Colors of nodes indicate species that will survive or go extinct after initialization. b) The example food web after initialization and before fishing effort (t=4000). Colors of nodes indicate species that will survive or go extinct after fishing. Indices (i) – (iv) indicate the order of extinctions during simulation. c) The final structure of the food web after the fixed harvesting effort (t=8000). d) Example simulation of a FE treatment simulation with effort set at 20. e) Example simulation of an Open Access treatment simulation.

We incorporated two economic models driving fishing effort into the ATN’s ecological network structure with fisheries functioning as an additional node in the networks. After an initialization period of 4000 time steps, roughly 11 years in model time (see Methods; 32) (Fig 1b), each “conserved” food-web (see Methods; 32) is subjected to two fishery treatments (Fig 1c): (1) Fixed Effort and (2) Open Access. The Fixed Effort treatment uses fixed levels of fishing effort starting immediately after the initialization period and do not change within simulations (Fig 1d). In the Open Access treatment, on the other hand, fishing Effort is unregulated (33). Instead, Effort adjusts in response to fishing profits, with Effort growth and decline occurring in response to positive and negative net profits respectively (Fig 1e). Profits are influenced by yield and market price. Market price is related to yield through a dynamic linear pricing model (see Methods; 32) (Fig S1). The Fixed Effort treatment serves as a control to the dynamics of Open Access fisheries, but it also has real world representations in strictly permitted fisheries used for subsistence fishing (34). The Open Access treatments simulate fisheries from their initialization at t=4000 and therefore start from a low initial Effort of 1. Effects of different economic conditions are studied by parameter sweeps across levels of price sensitivity to yield (*b*), Effort’s sensitivity to changes in profit (*μ*), and maximum price (*a*) paid for the harvested species. See Methods section for more (32). Fisheries are single-species, with the harvested fish per simulation labeled, *H*.

We use our dynamic model to evaluate the economic and ecological factors which determine: (1) the impact of fishing on harvested and non-harvested species (ecological impacts), (2) the conditions for fishery “success” (i.e., a sustained non-0 fishing effort), and (3) the different ecological impacts of fisheries within Fixed and Open Access regimes.

When we implement fishing through the Fixed Effort treatment, higher Effort levels increase *H* mortality, intuitively causing more biomass depletion (Fig S2), more extinctions (Fig S3), and quicker times to extinction (Fig 2a) for the harvested species, *H*. Among the hundreds of ecological factors analyzed (see Methods), we found those *H* extinctions to be more prevalent at higher trophic levels (Fig S4; Fig 2b), reflecting numerous empirical examples (35-37). Interestingly though, the best ecological predictor of *H*-extinctions, the population biomass of *H* at the start of fishing (*B*_*H*0_; Table S1), has a non-linear effect (Fig 2b; Fig S5a). Compared to the lowest *B*_*H*0_, moderate increases in starting population biomass decreased *H* extinction prevalence, as more abundant harvested populations are resistant to the extraction induced mortality. However, for all but the highest trophic levels, we saw that further increases in *B*_*H*0_ escalate extinction risk of *H*.

**Figure 2:**
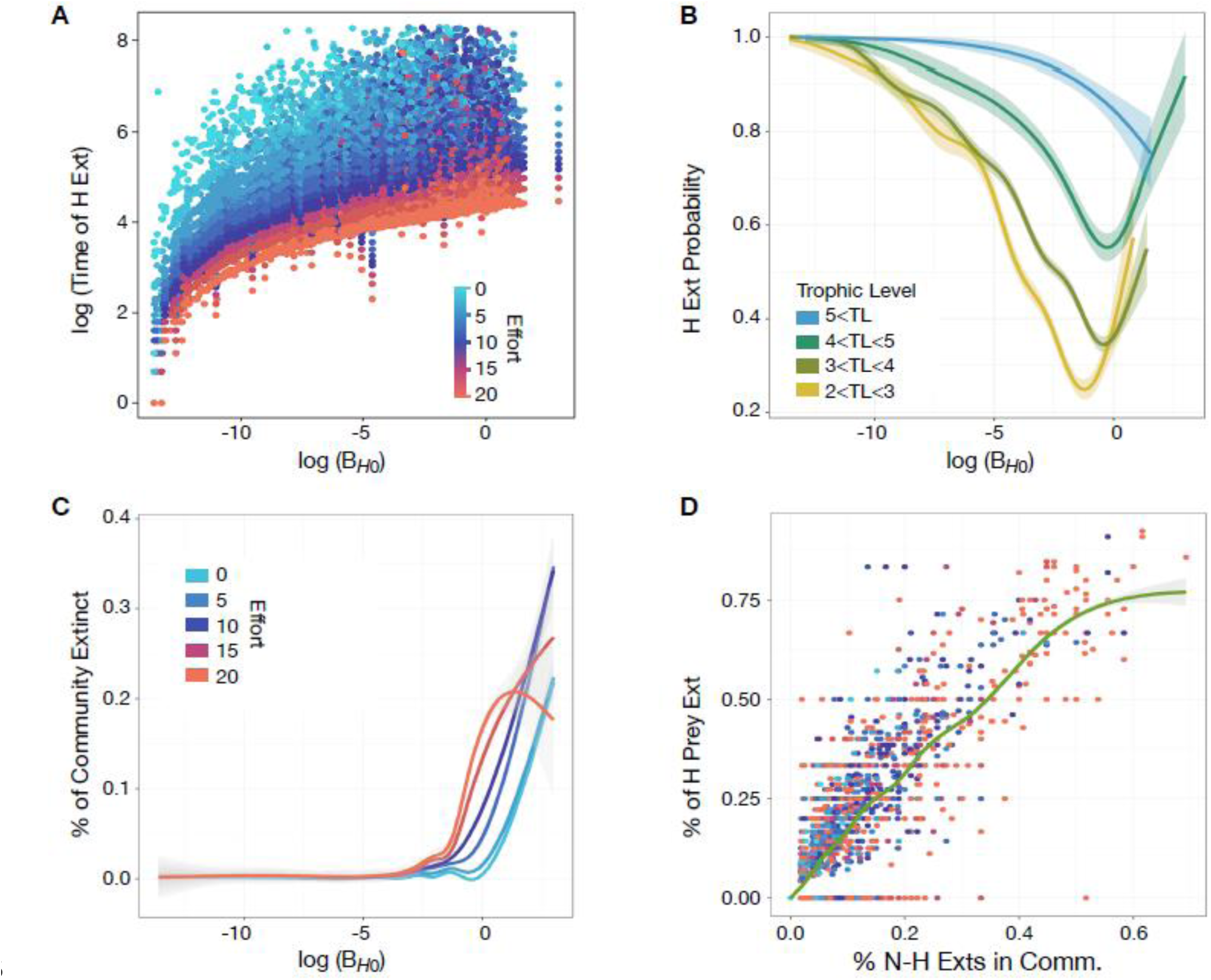
Fixed Effort results. a) Scatterplot showing the effect of log(*B*_*H*0_) on time to *H* extinction. b) Generalized additive model (GAM) binomial regression of the effects of log(*B*_*H*0_) on the probability of fishing resulting in *H* extinction across four trophic level groupings of *H*. Lines represent fit and shaded regions show 95% confidence interval. Including log(*B*_*H*0_), trophic level of *H*, and Fixed Effort level gives the binomial GAM results: 51.1% of variance exampled at *R*^2^=0.57, UBRE=-0.32. c) GAM regressions showing the percent of the community lost based on log(*B*_*H*0_). Lines represent fit and shaded regions show 95% confidence interval. Higher Effort levels (e.g. E=15, 20) do present a non-linear trend at high *B*_*H*0_. This is due to the expedited *H* extinctions at high Effort levels ending the active fishing disturbance on the rest of the community quicker than lower Effort levels. See Fig S6 for more. d) Scatter plot showing positive correlative relationship between the percent of community lost to *N-H* extinctions and the percent of *H*’s prey items lost. Colors represent Fixed Effort values similar to Fig 2a and the green line represents GAM regression fit with 95% confidence interval.

This non-linearity occurs because fishing higher *B*_*H*0_ generally induces greater levels of variability in the rest of the community’s populations (Fig S6; see Methods), mirroring past work that finds greater population variability when removing species with higher biomasses (29) (Table S2). Higher levels of variability generate extinction cascades of non-harvested species (*N-H* extinctions; Fig S7) which threaten *H* through its prey items. The association of *B*_*H*0_, population variability, and *N-H* extinctions functions as the mechanism behind *B*_*H*0_ causing higher levels of *N-H* extinctions (Fig 2c: Fig S5b; Fig S8). In fact, *B*_*H*0_ is also the best single pre-fishing predictor of *N-H* extinctions (Table S3) and its effects are exacerbated when higher Effort levels induce stronger perturbations in the community (Fig 2c; Fig S6; Table S3). The majority of these *N-H* extinctions occur downstream from *H* (Fig S9a), and are trophically close to *H* (Fig S9b) with the average distance becoming closer with a higher number of trophic links to *H* (Fig S10). As such, we clearly see that the degree of losses in *H*’s prey options increases proportionally with general *N-H* extinctions (*β* = 1.5, *p* < .0001, *R*^2^ = 0.75; Fig 2d). The relatively high level of downstream extinctions relative to upstream extinctions is at least partially due to the size constraints on the food-webs and the fish trophic levels studied here, but these results do indicate that perturbations due to fishing can cycle through the food web and threaten even highly abundant harvested species through their prey items.

In the Open Access treatment, *B*_*H*0_ also played a critical role in fishery sustainability as it was a principal driver of market dynamics. Open Access Effort is dynamic, capable of both declines and growth (Fig 3a). Growth in Open Access Effort is a function of *B*_*H*0_ and the maximum price of *H*, the parameter *a*. Higher max price reflects higher base demand for *H* and higher *B*_*H*0_ provides potential yield to meet demand, thereby driving greater profits and Effort (Fig S11a). Dependent upon demand and yield, Effort levels can range from low to high values. However, with sufficiently low demand or yield, net profit is consistently negative, Effort declines to 0, meaning the fishery fails to sustain itself (Fig S12). Consequently, harvesting the most abundant fish per web sustained more fisheries (78% of simulations) than fishing randomly chosen fish populations (26% of simulations). Though, higher prices/demand can sustain Effort on low abundance species by supporting higher profits on lower yield (Fig S13). This growth potential in the combination of max price (*a*) and harvestable biomass (*B*_*H*0_) strongly predicts the peaks in Effort early in Open Access fisheries’ time series (Fig 3a&3b; Table S4; Table S5).

**Figure 3:**
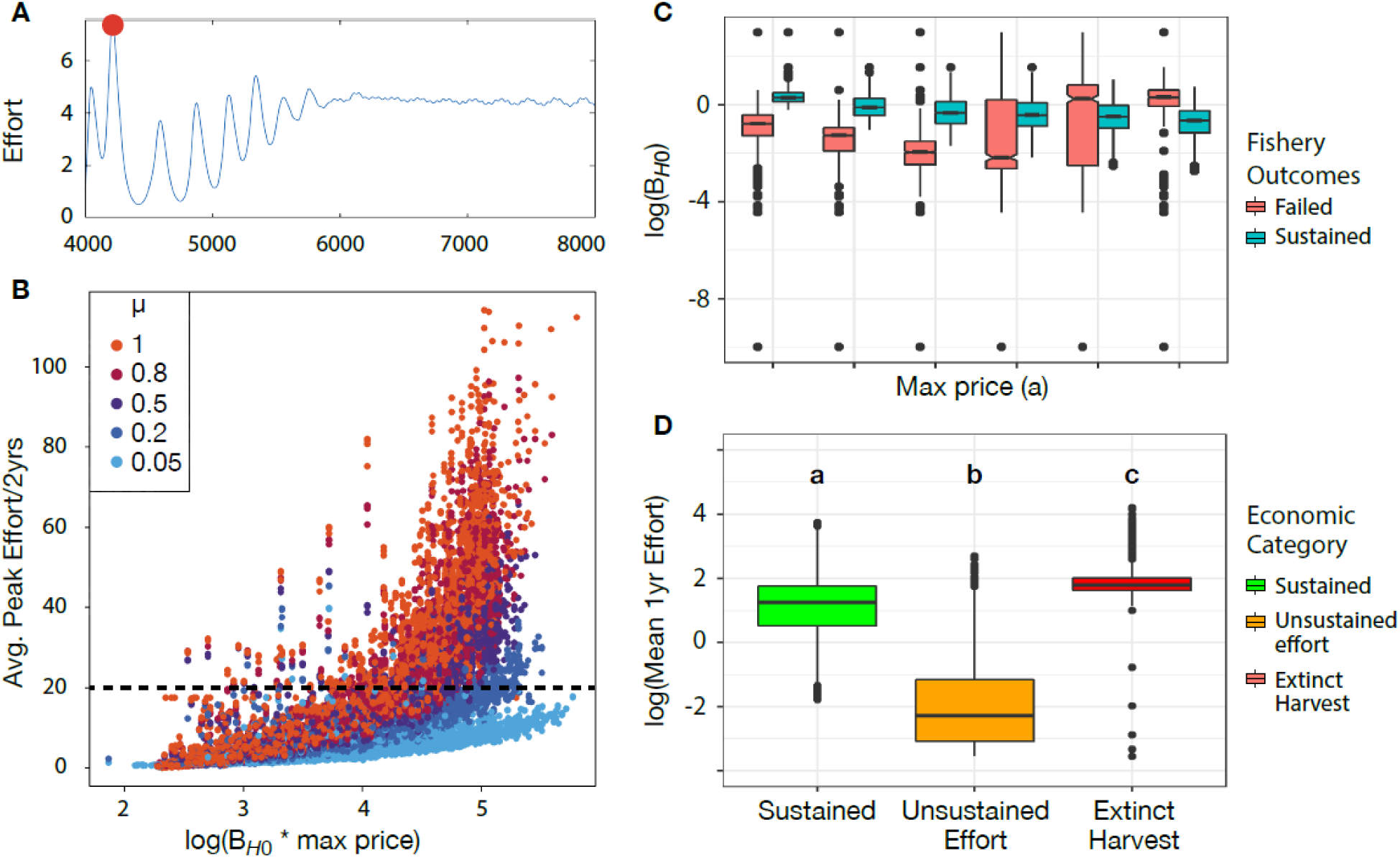
Open Access results. a) Example of effort dynamics where red dot represents a local peak effort. b) Peak efforts averaged across two years per the product of starting biomasses of harvested species (*B*_*H*0_) and max price (*a*). Colors indicate the value of *μ*, economic sensitivity/reactivity of the Effort per simulation. c) Box-plot showing *B*_*H*0_ of failed and sustained fisheries across different Max Price (*a*) when fishing the max *B*_*H*0_ fish population per food-web. Increasing *a* supports sustained fishing effort in less abundant fisheries until spiking effort peaks produced by higher profits when fishing abundant species cause the *H*-extinctions and fishery failure. d) Three economic outcomes characterized by efforts averaged across the first fishing year. Sustained (green) fisheries exhibit intermediate efforts producing enough profit without collapsing the resource. Unsustained (orange) fisheries failed before causing species extinctions because their profits were lower than harvesting costs. Extinct (red) fisheries failed because high profits produce high effort peaks causing the harvested-species extinction. Significant differences indicated by different letters (Tukey HSD, *p* < 2*e*^−16^). In boxplots of Fig 3c and 3d, boxes represent the interquartile range with the horizontal line showing the median, the lower box representing the 25 percentile, and the upper box showing the 75 percentile. Upper and lower lines extending from the boxes show the most extreme values within 1.5 times the 75^th^ and 25^th^ percentile respectively. Outliers are shown as single dots.

However, growth itself, if unchecked, can also lead to Effort declines during the “cycle of adjustment (38),” an empirically detected bio-economic process (39). That is, we see Effort reductions (Fig 3a) when past fishing effort has either over-supplied and caused market-saturation (reductions through price sensitivity to yield, *b*) or over-fished *H* (reductions through a lack of biomass from which to profit). These mechanisms of reactionary Effort reduction can function as “self-corrections” that potentially protect against excessive fishing Effort, allowing for regrowth in the harvested populations. These self-corrections, coupled with the potential for Open Access fisheries to fail to sustain Effort at low *B*_*H*0_, explain the relative lack of *H* extinctions in the Open Access tests (Fig S11b). However, especially in fisheries with high growth potential (high prices and *B*_*H*0_) where Effort is highly sensitive to profit (high *μ*), the combination of sufficiently high price (*a*) and high *B*_*H*0_ elevates Effort to levels (Fig 3b) inducing *H* extinctions before self-correction occurs (Fig 3c). Analyzing Open Access fishery outcomes reveals three general economic categories: (1) failed fisheries due to low *B*_*H*0_ and price (*a*) failing to maintain or grow Effort, (2) failed fisheries due to high *B*_*H*0_ and price driving *H* extinctions through excessive Effort, and (3) sustained fisheries existing in a middle ground between the two (Fig 3d).

These three general economic outcomes underpin much of the relationship between the harvested species and fishing Effort in Open Access fisheries. In particular, these results indicate that *H* extinctions were actually most common when harvesting highly abundant *H* (species with high *B*_*H*0_) as unregulated growth in Effort depleted fish reserves (Fig S14). This also formed the basis of the non-linear effect *B*_*H*0_ had on Open Access fishery persistence (Fig 4a) and the level of Effort sustained by the end of the simulation (Fig 4b). Additionally, while *H* extinctions were relatively less common, the link between harvestable biomass and Effort growth reversed the relationship between *B*_*H*0_ and time to *H* extinction seen in Fixed Effort fisheries (compare Fig 2a to Fig 4c). Finally, the dynamics of Effort in response to yield also drive the different effects of fishing on the rest of the community between Open Access and Fixed Effort fisheries.

**Figure 4:**
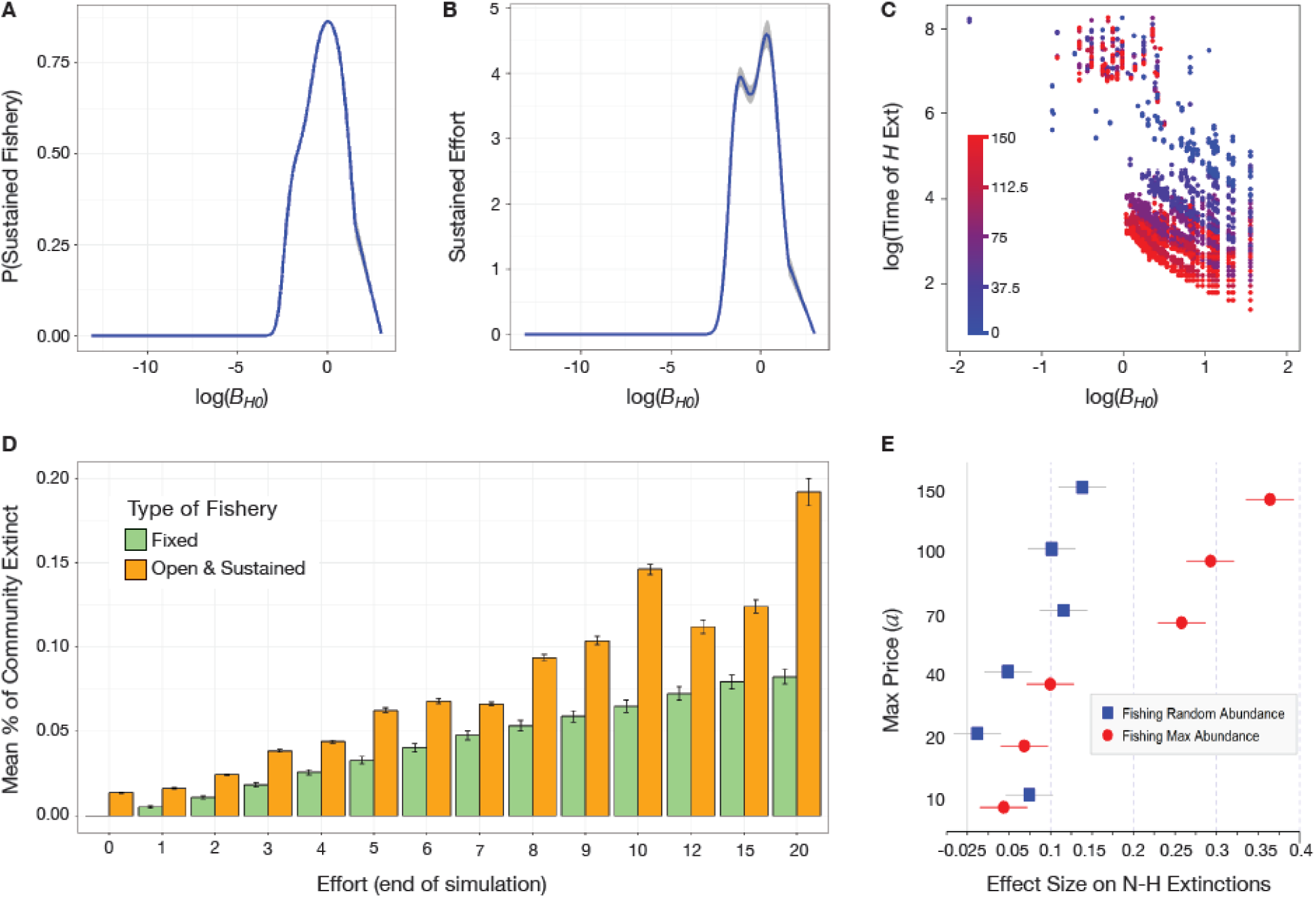
Open Access fishery’s economic and ecological sustatinability results. a) Probability of producing a sustainable Open Access fishery across the *B*_*H*0_ of *H*. Graph depicts the non-linear output of a binomial GAM regression with 95% confidence interval (*R*^2^ = 0.47^* **^, UBRE=-0.20). b) Sustained Open Access effort measured as average of final 400 time steps approaching t=8000. Graph depicts non-linear output of a gaussian GAM regression with 95% confidence interval (*R*^2^= 0.37, GCV=9.60). c) Scatterplot showing the effect of log(*B*_*H*_) on time to *H* extinction in Open Access simulations. Note the qualitatively reversed relationship from Fixed Effort fisheries. Color gradient represents the max price effort (*a*)* the market reactivity (*μ*). d) Bar graph with standard error comparing the percent of the community lost through *N-H* extinctions in Fixed Effort fisheries to Open Access fisheries with comparable effort levels at t=8000. E) Hedge’s G effect size comparison of the two fishery treatments (Fixed and Open Access) on *N-H* extinctions across different max prices (*a*). Dots represent effect size and lines show 95% confidence intervals. Comparisons are made within webs at comparable effort levels (see Methods; 32).

The profit-driven growth in Effort caused by high *B*_*H*0_ means that (given sufficient demand) economic factors incentivize subjecting food-webs to high levels of harvesting pressure on their more abundant species. This induces more variability in the biomass of the rest of the community than Fixed Effort fisheries (Fig S15). Expectedly (Fig S7), market generated variability induced more non-harvested (*N-H*) extinctions than Fixed Effort simulations. This was the case whether comparisons between Fixed Effort and Open Access results were made with attainable Efforts levels across all webs (Fig 4d; Fig S16; Fig S17) or strictly within each web at the most comparable Effort levels between fishery treatments (Fig 4e).

While our model framework has not yet been applied to multi-species fisheries or to realistically regulated fisheries, it can qualitatively reproduce past theoretical results (29) (Table S2), well-documented empirical patterns (Fig S4), and output from models specifically trained on empirical data (23) (Fig S18). This gives us confidence in vetting our results and further applying this framework towards the study of operational ecosystem based fisheries management (EBFM) strategies (27). Model output demonstrates the striking nonlinear effects of the population biomass of harvested species (*B*_*H*0_) on both the ecological and economic sustainability of fishing. It also reveals the role of *B*_*H*0_ in driving variations in Effort and exacerbating the ecological costs of harvesting in Open Access fisheries. Temporary fluctuations in Effort above sustainable levels are a known process in the cycles of adjustment seen in Open Access fisheries (33), but we show here that the indirect effects of those fluctuations can drastically change food-web structure. Overall, our results indicate that considering the sustainability of even one aspect of an ecosystem service likely requires a holistic accounting of both the ecology that provides the service and the economy that the service provides.

## Supporting information

Table S6 - Descriptions of Factors

## Acknowledgements

We would like to thank Kailin Kroetz, Maria Isidora Avila, the Valdovinos lab, and the researchers at the “Socio-Environmental Networks of Common Pool Resources” Working Group at National Socio-Environmental Synthesis Center (SESYNC) for feedback on study design, results, and interdisciplinary communication. We would also like to thank John Megahan for his assistance in producing the figures for publication.

## Funding

No outside funding was applied to this study.

## Author Contributions

FSV conceived study. FSV, PG, and VC designed study. VC and PG wrote and implemented simulation code. PG and FSV completed analysis and wrote manuscript.

## Competing Interests

The authors declare no competing interests.

## Data and Materials Availability

Simulation code and data will be available upon acceptance at the repository https://github.com/fsvaldovinos/Bioecon_Fisheries.

## Supplementary Material

Materials and Methods

Figures S1 – S18

Tables S1 – S6

References (40 – 53)

