## Supplementary Figures and Tables for "Integrating Economic dynamics into Ecological Networks: The case of fishery sustainability"

**<u>This PDF file contains the following content:</u>**

**Materials and Methods**

**Figure S1:** Linear Pricing Model diagram.

**Figure S2:** Mean percent loss of biomass of the harvested species due to different Fixed Effort levels.

**Figure S3:** Fixed Effort ecological outcomes via harvested (*H*) extinctions and non-harvested (*N-H*) extinctions.

**Figure S4:** Effect of H trophic level on H extinction probability in Fixed Effort simulations.

**Figure S5:** Effect of $\log(B_H)$ on $H$ extinction probability (Fig S5a) and $\geq 1$ *N-H* extinction probability (Fig S5b) across all Fixed Effort simulations.

**Table S1:** Summary stats on *H*-extinction probability.

**Figure S6:** Relationship between $B_{H0}$ and the induced mean variability in the rest of the community across the initial 3 years of fishing.

**Table S2:** Summary stats on interaction strengthen analysis, similar to Berlow et al 2009.

**Figure S7:** Relationship between the 3 year mean community variability and percent of food web lost to N-H Extinctions.

**Figure S8:** Differences between mean $B_{H0}$ fishing efforts that cause ≥1 *N-H* extinction (red) and no *N-H* extinctions (black) across all tested Fixed Effort values.

**Table S2:** Summary stats on N-H extinction probability.

**Figure S9:** Summary of locations of *N-H* extinctions in food-webs in Fixed Effort simulations.

**Figure S10:** The average distance away from $H$ of all *N-H* extinctions from the Fixed Effort simulations (left) and the Open Access simulations (right) as a function of the number of trophic links of $H$.

**Figure S11:** Effects of Open Access Max Price on effort and ecological outcomes.

**Figure S12:** Open Access sustainability across max price and fish choice treatments.

**Table S4:** Summary stats on peak Open Access Effort values w/in first year using linear model.

**Table S5:** Summary stats on peak Open Access Effort values w/in first year using generalized additive model.

**Figure S13:** Sustained and failed fisheries categorized by $B_{H0}$ across max price ($a$) values.

**Figure S14:** Open Access extinction probabilities across $B_{H0}$.

**Figure S15:** Hedge’s G comparison of the mean community stability of food webs fished through the Open Access management and the closest Fixed Effort value across the six max price values tested.

**Figure S16:** Comparisons of mean species lost to N-H extinctions between Open Access and Fixed Effort fisheries at comparable effort levels.

**Figure S17:** Bar graph comparing the number of *N-H* extinctions in Fixed Effort fisheries to Open Access fisheries with comparable effort levels at t=8000.

**Figure S18:** Comparison with results from Smith et al 2011.

Other Supplementary Material for this manuscript include:

**Table S6:** A description of ecological and economic factors used in the analysis of each fishing treatment.

**Materials and Methods**

Experimental Design

The objective of this study was to expand ecological network theory to include dynamic economic components. This Materials and Methods section describes the creation of ecological-economic networks and how they were analyzed. The ecological network models are composed of two major parts, the underlying structure of the network of interactions between species and the population dynamics of the interacting species (30, 40). In our model framework, we expand upon preliminary work (11) and add economic dynamics to the established population dynamics.

Network/Food Web Structure

Network structure describes the trophic links between resource, prey, and predator populations irrespective of the strength of the link. Initially, 1100 trophic food webs were created as networks using the Niche Model (31). All food webs contained 30 trophic species with a connectance of 0.15 (29) within an error of 0.025 (31). Trophic species ($S$) define a population of individuals with similar resources and consumers. Connectance ($C$) is the fraction of realized trophic links ($\frac{L}{S^2}$) where $L$ is the actual number of realized links. Trophic species are assigned a niche value across a single trait axis (considered body size in this case, see Methods: Ecological and Economic Dynamics section below) defined by a feeding range and center along the trait axis. Species $P$ eats species $V$ if $V$'s niche value lies within $P$'s consumption range. This process does allow for the species' consumption range to be at a higher niche value which allows for cannibalism. The consumption range of the lowest species in the niche axis is fixed to 0 assuming at least one primary producer/basal species. Each of the 30 trophic species is assigned values iteratively until 1) the web is connected (can't be divided into two independent webs, 2) every consumer species is linked to at least one basal species through a trophic chain, and 3) the realized connectance is within the error range set by $C$. Trophic species that can be potentially harvested are labeled 'fish' for ease of discussion. The fish species are chosen among the consumer species of each web with a Bernoulli's trial ($p = 0.6$) (11). Niche Model food webs have been shown to exhibit empirically observed patterns in field webs (distribution of trophic species across different trophic levels, mean trophic chain length, etc.), especially in aquatic systems (31, 41, 42).

Table 1: Parameter/Function definitions, values, and sources. Symbol descriptions: *: carnivore species, +: herbivore species, $: invertebrate species, =: fish species.

| **Ecological Parameter/Equations** | **Description** | **Value** | **Source** |
|---|---|---|---|
| $K$ | Carrying capacity | 1 | Brose *et al.* 2006 (43) |
| $e$ | Assimilation efficiency | $0.85^{*}/0.45^{+}$ | |
| $a_x/a_r$ | Allometric constant | $0.314^{\$}/0.88^{=}$ | |
| $y$ | Maximum consumption rate | $8^{\$}/4^{=}$ | |
| $Z$ | Body size ratio | 100 | |
| $B_0$ | Half-saturation biomass | 0.5 | |
| $h$ | Hill coefficient | 1.2 | Williams 2008 (42) |
| $B_{ext}$ | Extinction threshold biomass | $10^{-6}$ | Schneider *et al.* 2016 (44) |
| $\omega_{ij}$ | Preference of $i$ for $j$ | variable | N/A |
| $F_{ij}$ | Functional Response | variable | N/A |
| **Economic Parameters** | **Description** | **Value** | **Source** |
| $q$ | Catchability coefficient | 0.01 | Martinez *et al.* 2012 (11) |
| $c_o$ | Cost per unit Harvesting Effort | 0.01 | N/A |

Ecological and Economic Dynamics

Population dynamics of the trophic species on the food web are regulated through the allometric trophic scaling (30) extended to a multi-species food web to function as an Allometric Trophic Network (43, 45) (ATN). The ATN models the biomass ($\frac{\mu g C}{L}$) dynamics of trophic species through a system of consumer-resource equations (1) and (2).

$$\overbrace{\frac{dB_i}{dt}}^{\textbf{Producer Biomass}} = \overbrace{r_i(1 - \frac{\sum_{k \in producers} B_k}{K}}^{net\ primary\ production} - \overbrace{\sum_{j \in predators} x_j \frac{y_j}{e_{ji}} F_{ji} B_j}^{loss\ by\ predation} \tag{1}$$

$$\overbrace{\frac{dB_i}{dt}}^{\textbf{Consumer Biomass}} = - \overbrace{x_i B_i}^{maintenance\ loss} + \overbrace{\sum_{j \in prey} x_i y_i F_{ij} B_i}^{gains\ by\ predation} - \overbrace{\sum_{j \in predators} x_j \frac{y_j}{e_{ji}} F_{ji} B_j}^{loss\ by\ predation} - \overbrace{q_i E_i B_i}^{loss\ by\ harvesting} \tag{2}$$

Parameter values shown in equations (1) and (2) are described in Table 1. The functional response of consumers to prey biomass is given in equation (3). Note, $B_{0,j}$ denotes the half saturation biomass of species $j$ and that $\omega_{ij}$ denotes the preference of consumer $i$ for prey species $j$. Each preference, $\omega_{ij}$, equals the inverse of the total number of $i$'s prey species and changes through time when prey go extinct.

$$F_{ij} = \frac{\omega_{ij} B_j^h}{B_{0,j}^h + \sum_{k \in prey\ species} \omega_{ik} B_k^h} \tag{3}$$

The applicability of the ATN framework to models with such a large number of species stems from the ability to parametrize the physiological rates through a negative-quarter power law relationship with species' body masses, the single trait axis described in Network Structure section (30, 46). This is realized through the three rates reproduction, $R$, metabolism, $X$, maximum consumption, $Y$:

$$R_p = a_r M_p^{-0.25} \tag{4}$$

$$X_C = a_x M_C^{-0.25} \tag{5}$$

$$Y_C = a_y M_C^{-0.25} \tag{6}$$

where $a_r$, $a_x$, and $a_y$ are allometric constants which determine rates based on the body size $M_i$. The subscripts $C$ and $P$ indicate consumer and producer parameters respectively (30). The time scale of each food-web is set by setting the mass-specific growth rate of the basal species to one. With this as a reference and the assumption that basal species share the same body size, the mass-specific metabolic rates of all species are normalized by the time scale ($R_{ref}$), and the maximum consumption rates are normalized by the metabolic rates:

$$r_i = \frac{R_i}{R_{ref}} = 1 \tag{7}$$

$$x_i = \frac{X_i}{R_{ref}} = \frac{a_x}{a_r}\left(\frac{M_i}{M_{ref}}\right)^{-0.25} \tag{8}$$

$$y_i = \frac{Y_i}{X_i} = \frac{a_y}{a_x} \tag{9}$$

The body-size ratio between predators and prey ($\frac{X_i}{R_{ref}}$) is considered to be a constant ($Z$), a reasonable approximation in an aquatic system (43). This allows $x_i$ to be expressed as below:

$$x_i = \frac{a_x}{a_r}\left(Z^{T_i - 1}\right)^{-0.25} \tag{10}$$

Where $T_i$ is the prey averaged trophic level of species $i$ calculated from network topology (41). Model time is set similar to past work (47) where one model time step equals one real-time day.

Loss of a population’s biomass due to harvesting (labeled ‘loss by harvesting’ in equation (2) is measured as the rate of fishing Effort, $E_i$ equation (11), on species $i$ (48). Effort is a broadly applicable index used to measure the amount of fishing/harvesting taking place in a fishery, including capital and labor (33). Depending on the specific fishery, Effort can track the number of fishing lines, boats, workers, work hours dedicated to harvesting, etc. The results presented in this work would not qualitatively change based on the specific details of the Effort metric. This study focuses on single species fisheries. Therefore, $E_i = 0$ for all species that are not the harvested fish species, $H$. The fishing effort on $H$ is greater than or equal to 0 and changes in $E$ derive from the product of net profit and an adjustment parameter, $\mu$, representing the economic sensitivity of fishing effort to changes in net profit (48). The economic sensitivity of the fishery’s Effort ($\mu \in [0,1]$) describes the sensitivity of the Effort to changes in profit or loss. Net profit is defined as the product of price per unit of biomass harvested and the actual yield ($Y$) caught at the current $E$ level, equation (12), minus the costs per unit effort (48). The yield at any given time in the model is a product of the current level of effort, the available biomass of the harvested species ($H$), and $q$ the ‘catchability’ of $H$ per unit effort. Yield translates into supply and informs the market price, $p$, through the linear equation (13) were $a$ represents the maximum price and price sensitivity to yield is labeled $b$. Equation 13 is incorporated as a piece-wise equation such that $p = 0$ when $Y \geq \frac{1}{b}$. Finally, costs are removed from gross profit to reach net profit by subtracting $c_o E$ from $pY$ were $c_o$ is the fixed value of cost per unit effort. All Open Access fisheries start with $E(0) = 1$ to model a fishery from its initiation.

$$\overbrace{\frac{dE_i}{dt}}^{\substack{\textbf{Effort}\\\textbf{Level}}} = \overbrace{\mu}^{\substack{Sensitivity\\to\ net\ profit}} * \overbrace{\underbrace{pY}_{\substack{Gross\\Profit}} - \underbrace{c_o E_i}_{\substack{Costs\\per\ Effort}}}^{Net\ Profit} \tag{11}$$

$$Y = qEB_H \tag{12}$$

$$p = a(1 - bY) \tag{13}$$

<u>Experimental Setup and Treatment Design</u>

All 1100 food webs created with the Niche Model were randomly assigned initial biomass conditions per species using a uniform distribution, $U \in (0,1)$. The ATN framework shown in equations 1 and 2 was then used to simulate each food web for 4000 time steps in a fishing free stage (Effort=0). This initial fishing-free period limits possible effects of transient dynamics on results of fishery treatments. After the initial fishing-free period, only “conserved” webs which met fishery criteria are chosen to be subjected to the two different fishery treatments.

A food-web is considered conserved if :

(i) it is connected

(ii) every consumer species is linked to at least one basal species through a trophic chain

(iii) the number of remaining trophic species is higher than or equal to 20

(iv) with at least one fish species.

These criteria resulted in 480 unique conserved food webs which were subjected to an additional 4000 time steps of simulation in the fishing stage. Subjecting the initial food webs to these criteria eliminated the potential for analytical comparisons of fishery effects between categorically different types of food webs.

Fishing effort was applied as one of two fishery treatments, Fixed Effort and Open Access. In the Fixed Effort treatment, $\frac{dE}{dt} = 0$ and fishing effort is held constant from t=4000 to t=8000, during the entire fishing treatment. The tested fixed effort values are $E = [0,1,2,3,4,5,6,7,8,9,10,12,15,20]$. In the Open Access treatment, effort varies based on the interactions between economics and ecology through profit on yield described above. Open Access results are tested across the values of effort sensitivity to profit $\mu = [0.05, 0.2, 0.5, 0.8, 1.0]$, price sensitivity to yield $b = [0.01,0.05, 0.1, 0.5]$, and maximum price $a = [10,20,40,70,100,150]$. Initial effort in all Open Access simulations is set at 1 to model a fishery from its beginning.

Regardless of fishery treatment, fishing effort was applied to a single fish species per simulation, $H$, chosen at the end of the initial stage in conserved webs, at t=4000. We do not model by-catch or multiple-species fisheries. In the Fixed Effort treatments, every single fish was harvested. In the Open Access treatment, given the importance of population biomass seen in the Fixed Effort results, the identity of the harvested fish species was selected either 1) randomly across all available fish in the food web, labeled ‘Random’ in figures and results or 2) as the fish species with the highest biomass after the initial stage ($B_{H0}$ at t=4000), labeled ‘Max’ in figures and results. Webs with only one harvestable fish species were only used once per parameter combination per fishery treatment. These Open Access results could then be directly compared to their Fixed Effort counterparts.

All simulations were completed using Matlab 2018a and the solver ode45 for numerical integration (relative and absolute error tolerances both equal to $10^{-8}$).

Data Collection and Categorization

Ecological, economic, and network data was compiled initially at t=0, at the end of the initial fishing-free stage at t=4000, and at the end of the fishing stage t=8000. Species extinctions during simulations were considered at the threshold, $B_{ext}$ listed in Table 1 and are displayed in Fig S3 and Fig S11b. Beyond the time series data of each species in each food web simulation, a large number of variables describing pre- and post-fishing overall network attributes (e.g. connectance) and local network structure around the harvested species (e.g. number of direct trophic links) were also considered. A full listing of these variables appears in the Table S6.

Per simulation, 408 attributes were considered in both the Fixed Effort treatment and Open Access treatments. Additional variables were added to the Open Access treatment, for a total of 432, to detail the dynamics of effort through the simulation and to make direct comparisons to the Fixed Effort treatments (see Fig 4e). While effort levels were directly recorded in Open Access treatments, simulation results were also categorized as sustained or failed at the end of simulations. To avoid considering temporary troughs in effort as complete failures, effort levels were averaged over the last 400 time steps. Fisheries which did not maintain effort levels above 1% of starting effort values were labeled failures, while those that did were labeled sustained.

In each food-web simulation, dynamic variability of each population was measured using a population’s biomass time series’ coefficient of variance. A similar process was used to measure the variability in Effort in Open Access Fisheries. For the community’s dynamic variability, we used the mean of each population’s biomass time series’ coefficient of variance (C.V.) as a proxy. In other words, we took the C.V. of each species’ biomass time series and averaged them to get the mean variability of a community. This was done in the 400 time steps prior to the start of fishing, in the first year after the start of fishing, across the first 3 years after the start of fishing, and during the last year of fishing before each simulation ended. See Table S6 for full list of analyzed factors.

### Statistical Analysis

Given the large number of variables/factors, we used classification and regression tree (CART) analysis (49) through JMP Pro 13 to obtain variables of interest in the search for drivers of fishing induced changes across our webs and simulations. Once obtained, potential drivers were further explored using R 3.5.1 statistical software (50). Continuous variables were initially explored using generalized linear models. Binomial regressions were used for binary response data while gamma or Gaussian distributions were used for continuous response data. Models were trained and vetted on various resampled subsets of data to assess consistency of the model results and avoid the pull of outliers. In the case of biomass of the harvested species ($B_{H0}$) resampling indicated potential non-linear responses. As a result, we used Generalized Additive Models (GAMs) to account for and visualize $B_{H0}$’s effect on response variables using the mgcv package in R (51). When response variables were continuous and limited between 0 and 1 (Fig S7), beta regressions were used (beta package R) (52). Categorical variables can be incorporated into GAMs as factors (Table S3). In situations where GAMs were not appropriate, comparisons across categorical variables were done using Tukey’s HSD test (Fig 4d).

There are two major comparisons of ecological cost of harvesting through the Non-Harvested (*N-H*) extinction prevalence between the two fishery treatments. The first (Fig 4d, Fig S16, Fig S17) compares the average *N-H* extinctions associated with comparable efforts levels between the Fixed Effort and Open Access fishery treatments. Due to inherent variability in effort levels in Open Access treatments, not every fish in every web can be guaranteed to be fished at every effort level as was done in the Fixed Effort treatment. Therefore Open Access Effort per simulation was averaged across three time periods, the first year (Fig S16a), the third year (Fig S16b), and across the last year (Fig 4d) and then grouped into one unit Effort blocks. For example, 0< *Effort*<1, 1< *Effort* <2, 2< *Effort* <3, and so on along the tested Fixed Effort values. *N-H* extinctions from those Open Access simulations were then compared against these from the Fixed Effort simulation.

The first method’s focus on comparisons across effort requires comparisons across different webs. In order to constrain the *N-H* extinction assessment to within web comparisons, a second method was implemented. For each max price value ($a$), every single Open Access simulation’s *N-H* extinction level was compared to the *N-H* extinctions from the Fixed Effort treatment of the same fish in the same web with the closest matching Fixed Effort level when compared to the Open Access effort level averaged across the third year of Open Access fishing. The effect of fishery treatment was ascertained using Hedge’s G (effsize R package) (53) to compare effect size of fishing with Open Access with the Fixed Effort treatment as the control (Fig 4e). The same process was also used to compare other metrics, such as mean community biomass variability (Fig S15).

**Supplementary Figures and Tables:**

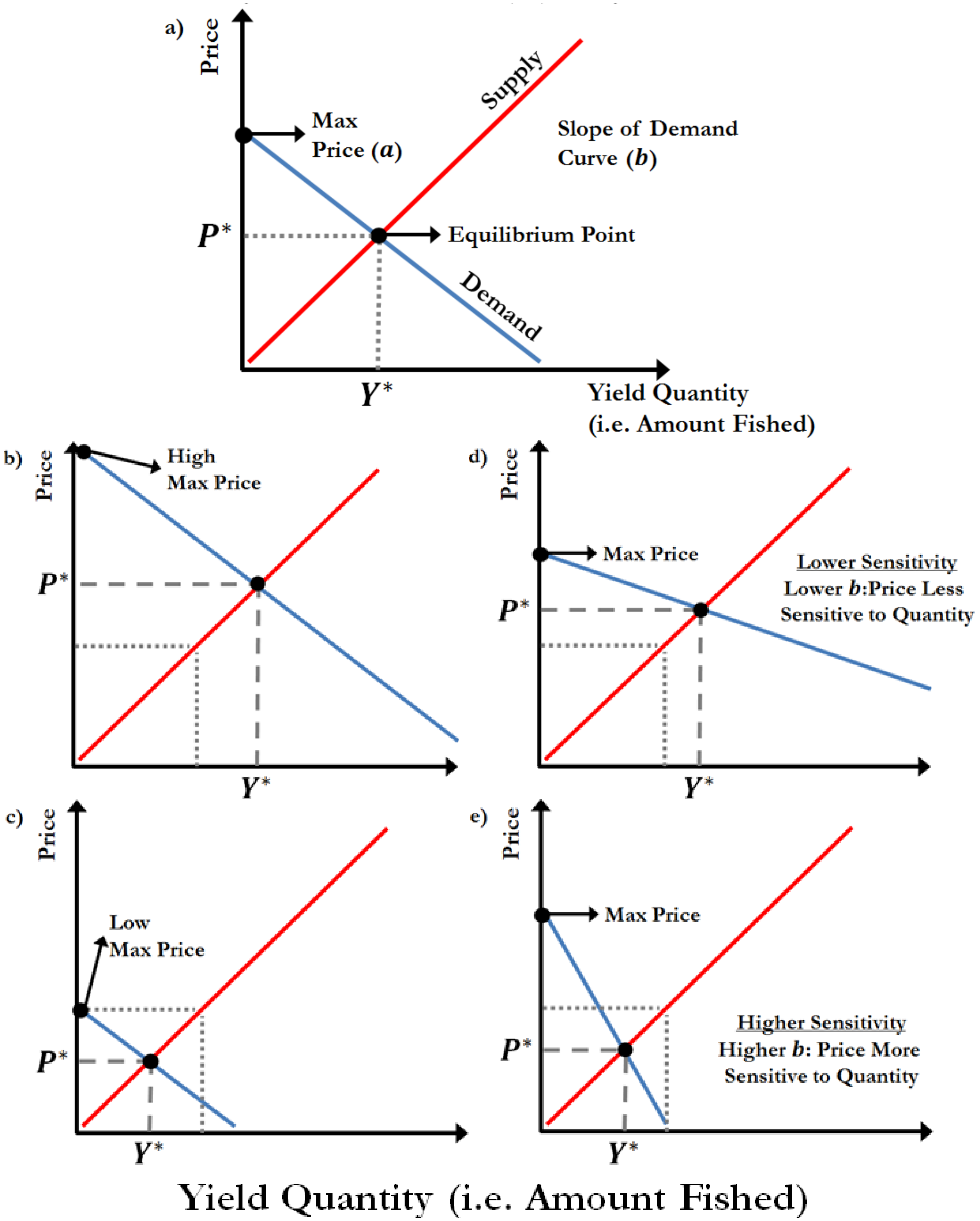


Figure S1: Linear Pricing Model diagram. a) Diagram representing the fundamental linear pricing model used in this study: $p = a(1 - b * Y)$. The red and blue curves represent the Demand and Supply curves respectively on the Price vs Yield Quantity axis. The intersection of the Supply and Demand curve represents the equilibrium where the rate of change of price and quantity supplied reaches 0. These values are denoted as $P^*$ and $Y^*$ for price and yield respectively. The maximum price the harvested species can take in the model is where the price curve intersects the y-axis as the supply goes to 0 and is labeled $a$. The slope of the Demand curve determines the sensitivity of the demand and price of the harvested species to changes in the quantity supplied/harvested. b) Example of raising the maximum price of the harvested species. This is akin to raising demand and subsequently increases the quantity supplied/harvested. c) Example of lowering the maximum price of the harvested species. This is akin to lowering demand and subsequently decreases the quantity supplied/harvested. d) Example of reducing the value $b$, the price sensitivity and slope of the Demand curve. This reduces the change in price/demand to in response to higher levels of Yield. e) Example of increasing the value $b$, the price sensitivity and slope of the Demand curve. This increases the change in price/demand to in response to higher levels of Yield.

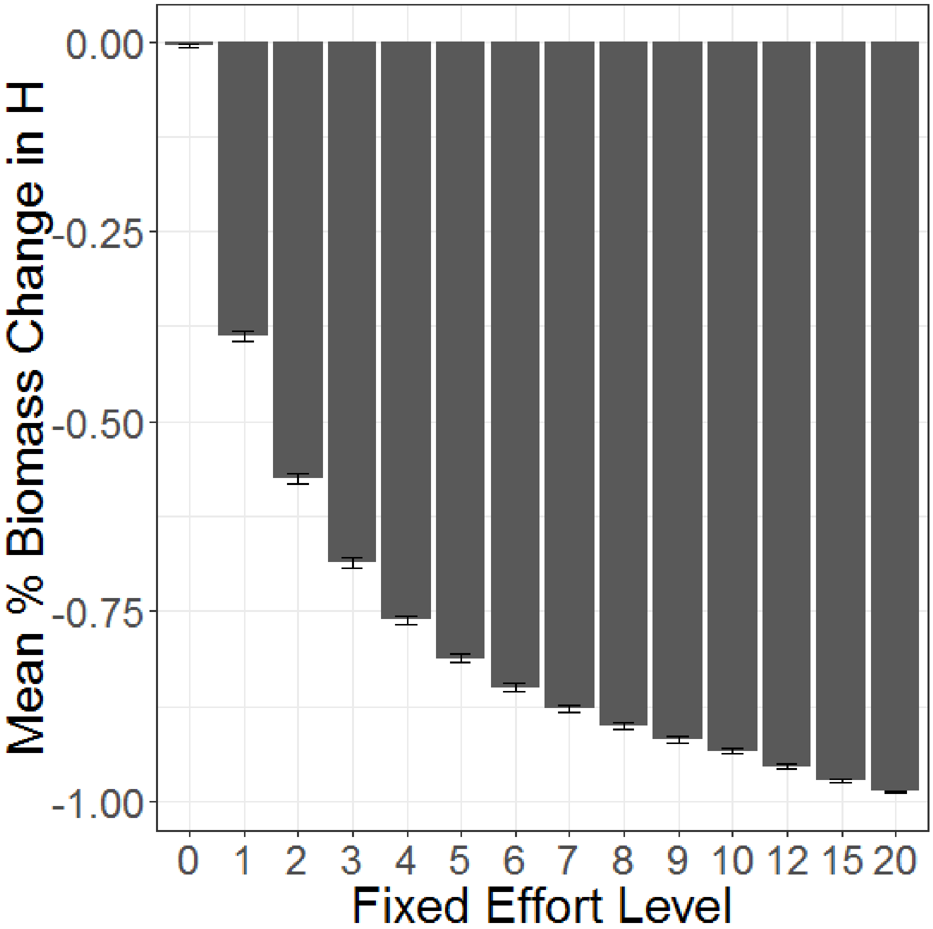


Figure S2: Mean percent loss of biomass of the harvested species due to different Fixed Effort levels. Error bars represent standard error.

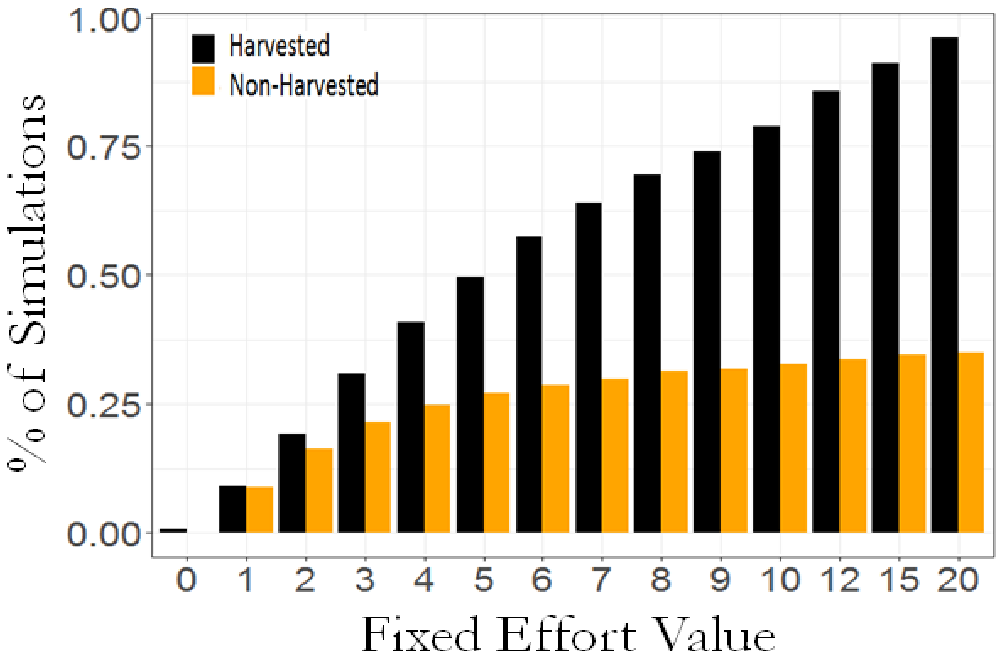


Figure S3: Fixed Effort ecological outcomes via harvested (*H*) extinctions and non-harvested (*N-H*) extinctions. A) Bar graph showing the percentage of all fishing simulations that resulted in *H* extinctions (black) and/or *N-H* extinctions (orange) at all tested Fixed Effort levels.

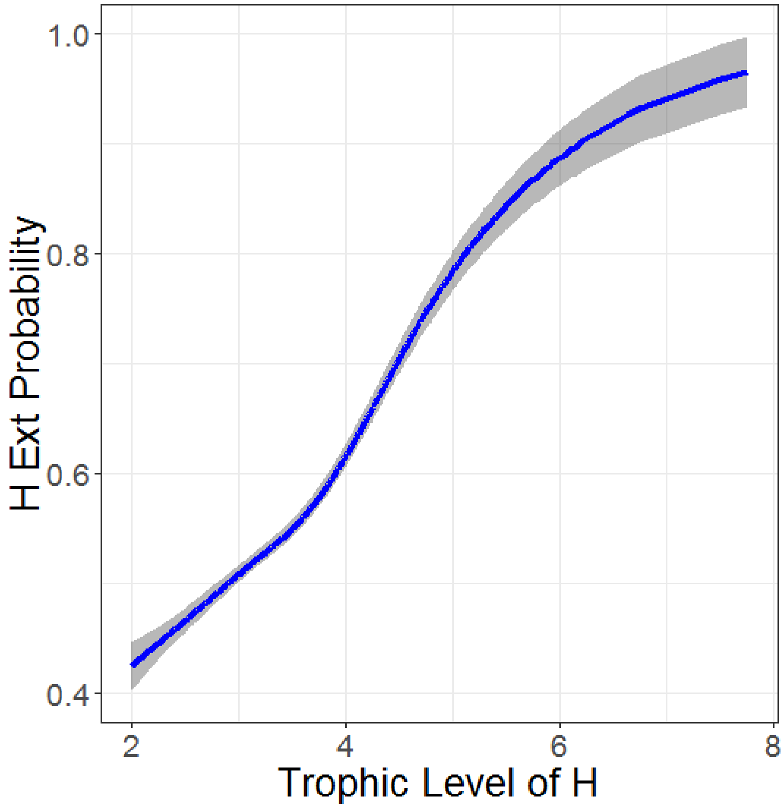


Figure S4: Line representing generalized additive model (GAM) regression with 95% confidence interval of trophic level of *H* alone versus *H*-extinction probability. $R^2$= 0.0251 Deviance explained = 1.94%, UBRE=0.35

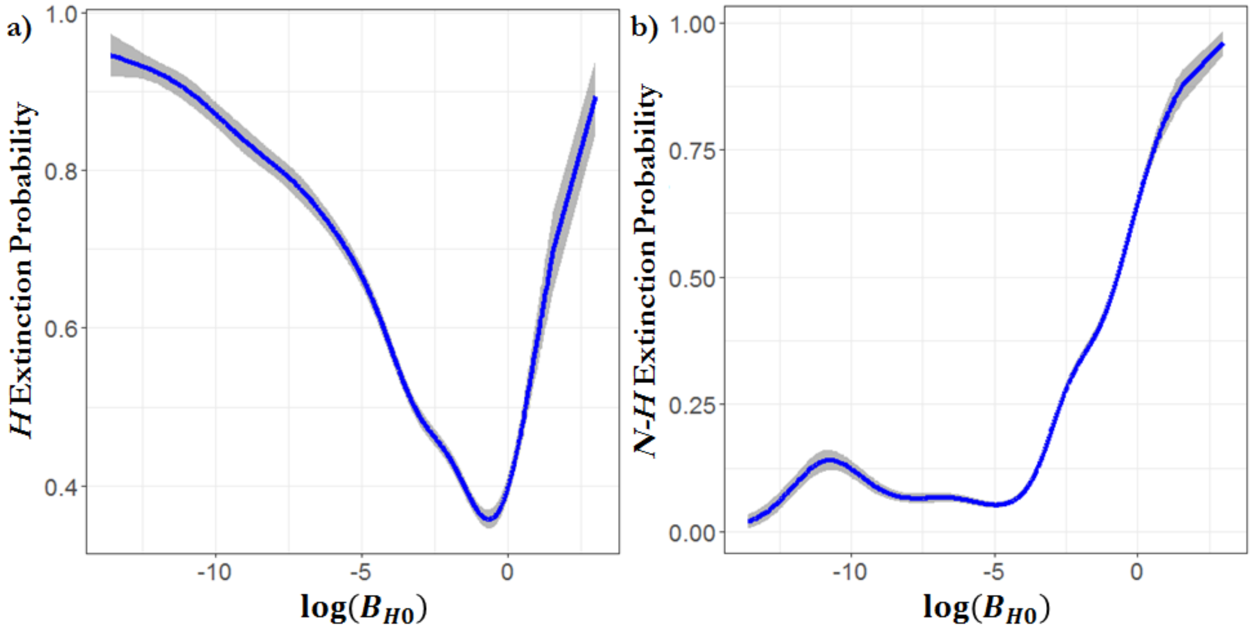


Figure S5: Cumulative effect of $\log(B_H)$ on $H$ extinction probability (Fig S5a) and $\geq 1$ $N$-$H$ extinction probability (Fig S5b) across all Fixed Effort simulations. Lines represent GAM regressions with 95% confidence intervals. Including Fixed Effort level in GAM regressions gives the following for the probability of H extinction: 44.7% of the variance with $R^2$=0.5, UBRE=-0.24. For the probability of $1 \geq NH$ extinction: 26.3% of the variance with $R^2$=0.29, GCV=-0.16.

Table S1: Summary results from binomial GAM regression on $H$-extinction probability as predicted by $\log(B_{H0})$, Fixed Effort level ($FE$), and the trophic level of $H$, $TL_H$. Graphs indicate effect across tested ranges where dotted region shows 95% confidence interval. Summary table provides summary statistics. Additional CART results are as follows: folded $R^2 = 0.69$, Highest contributors: Effort=46%, $\log(B_{H0})$ =11%.

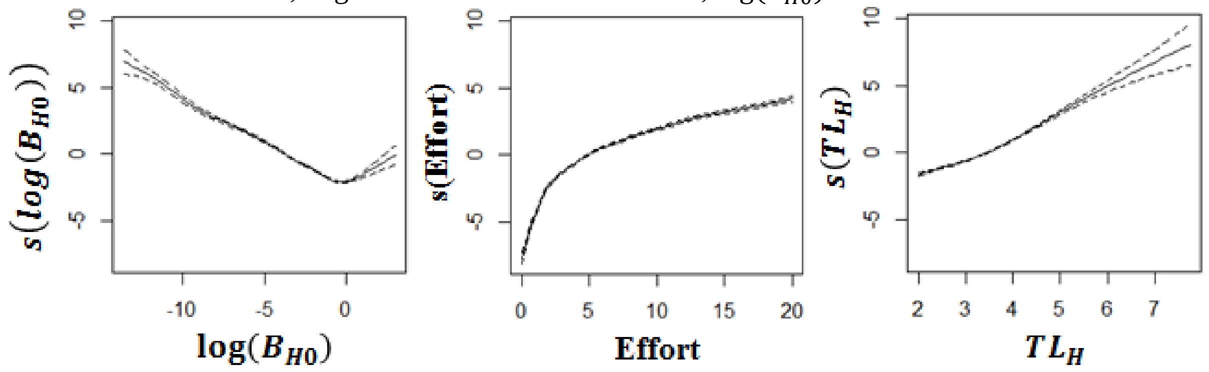


| Effect on Probability of H Extinction (Binomial GAM) | | | | | | |
|---|---|---|---|---|---|---|
| Model | p-value | UBRE | Scale Est | $R^2$ | Deviance Explained | AIC |
| $log(B_{H0})$ | $log(B_{H0})$**** | 0.26 | 1 | 0.11 | 8.28% | 48530.4 |
| $FE$ | $FE$**** | 0.36 | 1 | 0.36 | 30.4%% | 36835.15 |
| $TL_H$ | $TL_H$**** | 0.35 | 1 | 0.03 | 1.94% | 51869.99 |
| $log(B_{H0}) + FE + TL_H$ | $log(B_{H0})$**** $FE$****, $TL_H$**** | -0.32 | 1 | 0.57 | 51.1% | 25932.18 |

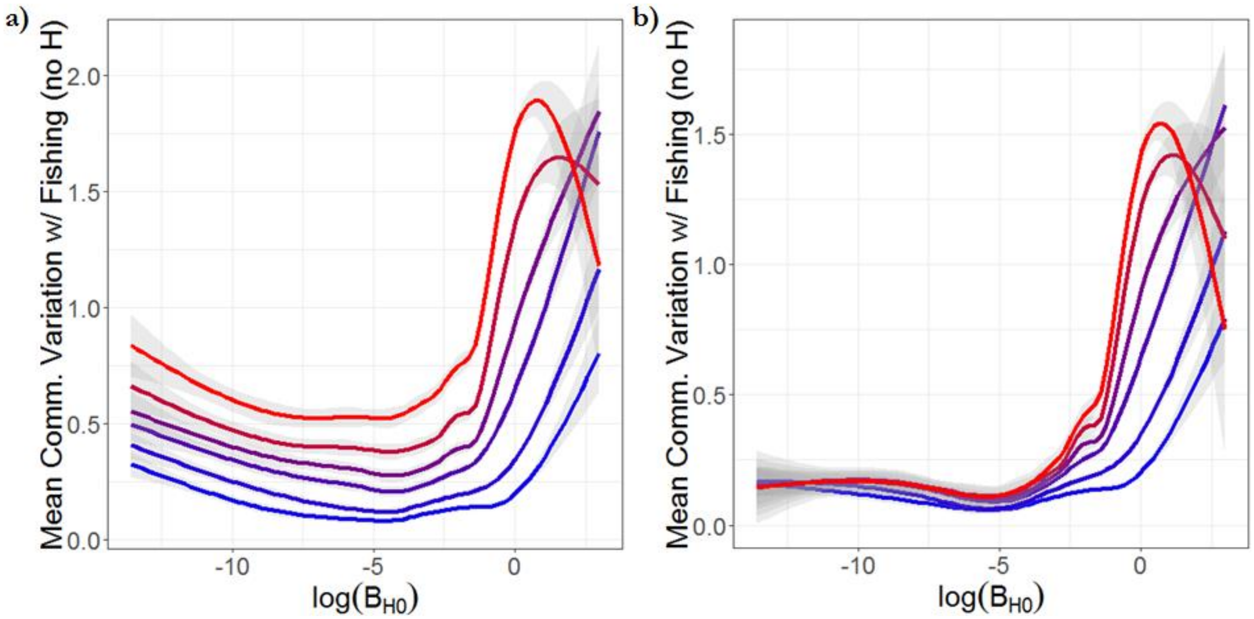


Figure S6: Relationship between the biomass of the harvested species ($B_{H0}$) and the induced mean variability in the rest of the community across the initial 3 years of fishing. Lines represent generalized additive model (GAM) regressions with shaded regions showing 95% confidence intervals and line colors represent a sample of Fixed Effort values from lowest (blue) to highest (red). a) The mean C.V. of each species with greater than zero biomass at the start of fishing. Coupled with Fixed Effort level, GAM analysis explains 50.1% of the variance with $R^2$=0.5, GCV=0.085, scale estimate=0.085 b) The mean C.V. of each species with greater than zero biomass at the start of fishing excluding the C.V. of the harvested species ($H$). Coupled with Fixed Effort level, GAM analysis explains 41.9% of the variance with $R^2$=0.42, GCV=0.091, scale estimate=0.091. In both regressions there is an eventual reduction in biomass variability at high levels of Fixed Effort due to a faster $H$ extinction time in the highest biomass $H$ populations. At such high $B_{H0}$, the faster $H$ goes extinct, the faster the active fishing disturbance ends, and the quicker the rest of the community can reach a new asymptotic behavior with less variability.

Table S2: Table of explanatory variables on the absolute magnitude of interaction strength between $H$ and other co-occurring species in each food web in each simulation. This is done to show our process can qualitatively approximate past results from Berlow et al 2009. Berlow et al 2009 removed a species $R$ from a food web and studied the effects on each remaining "target" species $T$. They defined interaction strength of the removal of $R$ as $I = B_T^+ - B_T^-$, where $B_T^+$ is the biomass of $T$ before removal of species $R$ and $B_T^-$ is the biomass of $T$ after $R$'s removal. Berlow et al 2009 results show the $\beta$ values of $\log_{10}(B_T)$ and $\log_{10}(B_{H0})$ to be 0.71 and 0.22 with $R^2$=0.65. The study design of Berlow et al 2009 removed all possible $R$ species from food webs at t=0 regardless of species identity and trophic level. While our study design only removed labeled fish species, thereby altering the number of species removals per web and their trophic level, results from our study better resemble those of Berlow et al 2009 as the Effort level rises, more rapidly depleting the harvested species, and more closely resembles the instantaneous species removal from their study design. Significance levels are all *** $p < 0.001$.

| Explanatory variables on $\log_{10}(\|I\|)$ | | | |
|---|---|---|---|
| **Effort** | $\log_{10}(B_T)$ | $\log_{10}(B_{H0})$ | $R^2$ |
| **1** | $\beta = 0.678^{***}$ | $\beta = 0.344^{***}$ | 0.5127 |
| **5** | $\beta = 0.680^{***}$ | $\beta = 0.238^{***}$ | 0.5331 |
| **10** | $\beta = 0.681^{***}$ | $\beta = 0.238^{***}$ | 0.5384 |
| **15** | $\beta = 0.683^{***}$ | $\beta = 0.242^{***}$ | 0.5427 |
| **20** | $\beta = 0.685^{***}$ | $\beta = 0.243^{***}$ | 0.5433 |

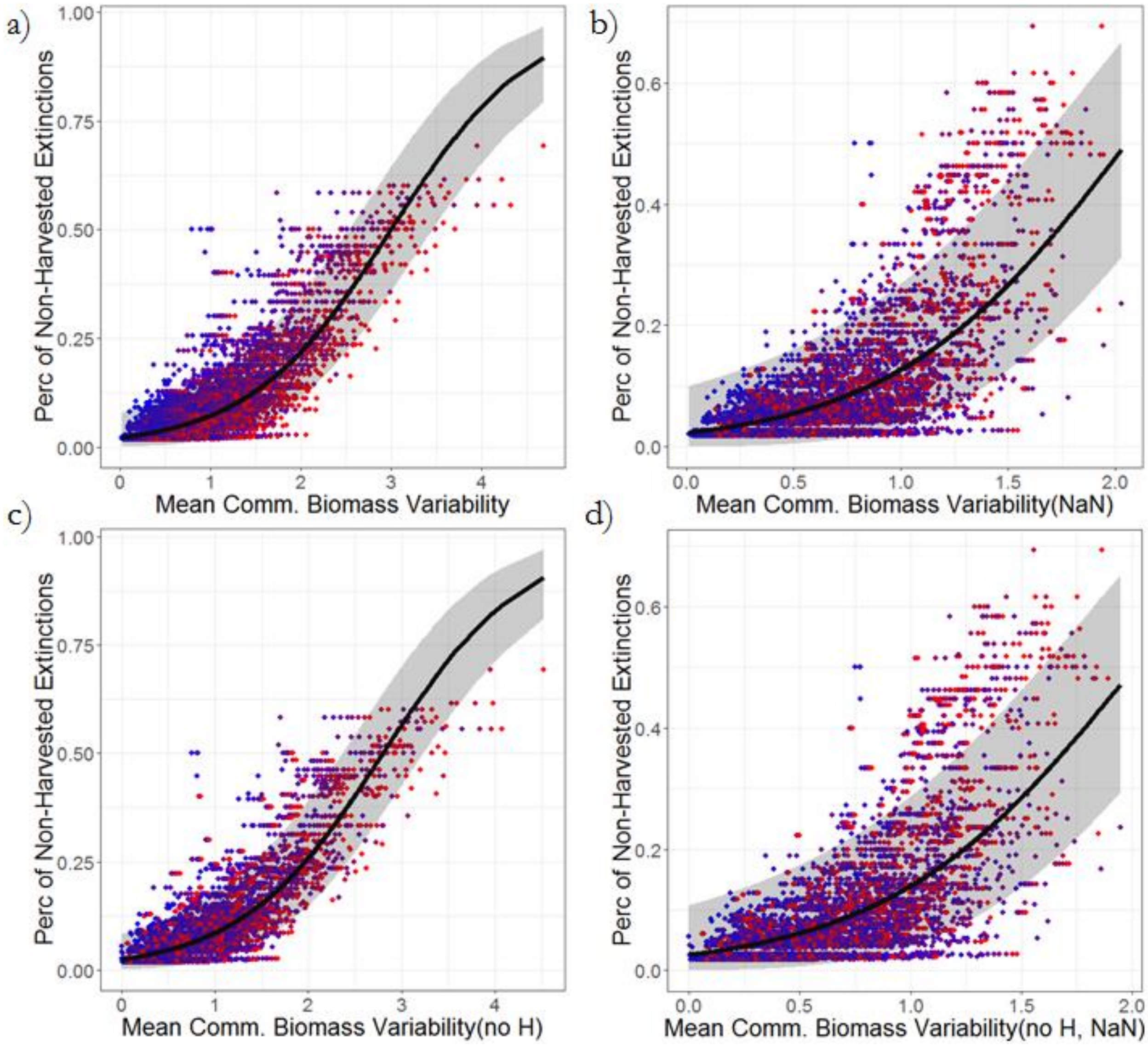


Figure S7: Scatter plots showing the correlations between mean community variability in the first 3 years of fishing and the percent of each food web lost to *N-H* extinctions in the Fixed Effort treatments. Dots colors represent the level of fixed effort. Lines in each panel depict a beta regression with 95% confidence intervals. Mean community variability is measured as the mean C.V. of the biomass time series of each species with greater than zero biomass at the start of fishing, with four variations to display the consistency of the result. a) Mean C.V. of community biomass including $H$. Beta regression results: $\beta = 1.28 \pm 0.007$, $z = 167.6$, $p < 2e^{-16}$, $\phi = 45.58$, $log - likelihood = 2.05e^{4}\ on\ 3\ DF$, $psuedo\ R^2 = 0.66$. b) Mean C.V. of community biomass including $H$, but not including any zeros in the time series of any species after it has gone extinct (zeros ignored in C.V. calculation). Beta regression results: $\beta = 1.83 \pm 0.016$, $z = 111.5$, $p < 2e^{-16}$, $\phi = 29.04$, $log - likelihood = 1.85e^{4}\ on\ 3\ DF$, $psuedo\ R^2 = 0.56$. c) Mean C.V. of community biomass not including $H$. Beta regression results: $\beta = 1.31 \pm 0.007$, $z = 175.8$, $p < 2e^{-16}$, $\phi = 48.86$, $log - likelihood = 2.09e^{4}\ on\ 3\ DF$, $psuedo\ R^2 = 0.69$. d) Mean C.V. of community biomass not including $H$, but not including any zeros in the time series of any species after it has gone extinct (zeros ignored in C.V. calculation). Beta regression results: $\beta = 1.80 \pm 0.016$, $z = 108.5$, $p < 2e^{-16}$, $\phi = 28.30$, $log - likelihood = 1.87e^{4}\ on\ 3\ DF$, $psuedo\ R^2 = 0.55$.

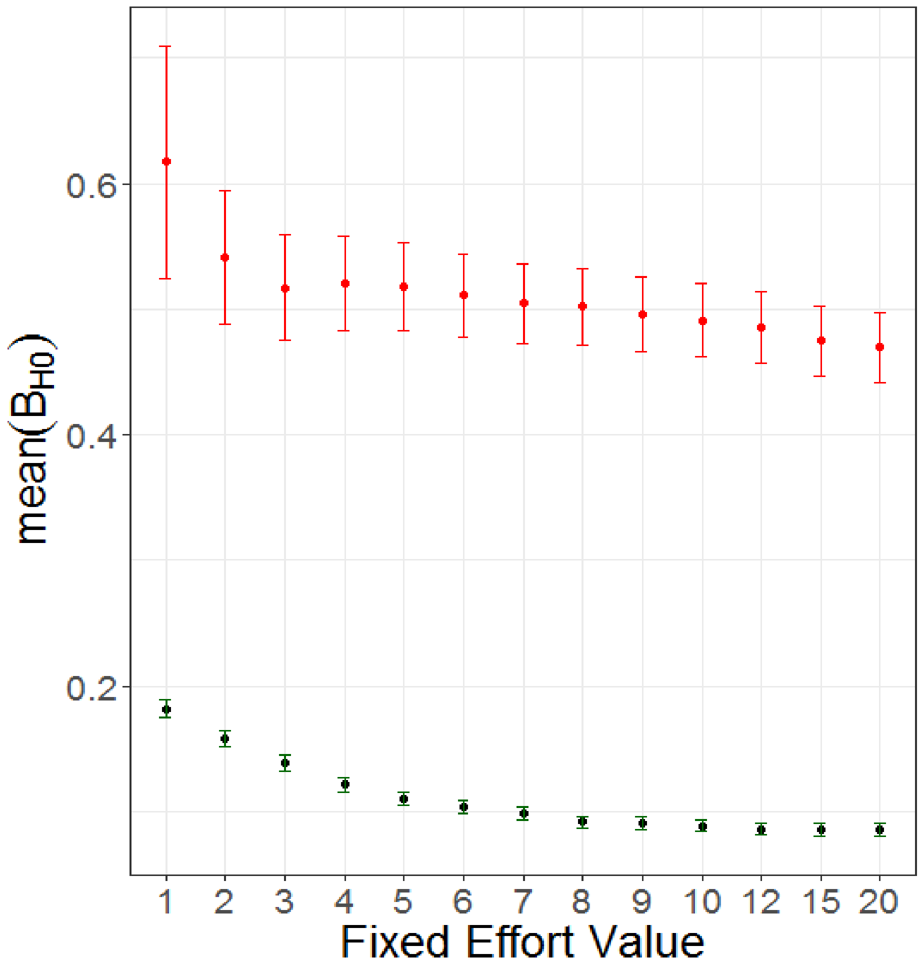


Figure S8: Differences between mean $B_{H0}$ fishing efforts that cause ≥1 *N-H* extinction (red) and no *N-H* extinctions (black) across all tested fixed effort values. The bars represent standard error. Standard deviations bars between each category do overlap but make the figure difficult to read in parts. For every fixed effort value, mean $B_{H0}$ between ≥1 *N-H* extinction (red) and no *N-H* extinctions (black) are significantly different for all Fixed Effort levels (t test: $< 2.2e - 16$ )

Table S2: Summary results from binomial generalized additive model (GAM) regressions on probability of at least one non-harvested extinction (*N-H* extinction) as predicted by $\log(B_{H0})$ and Fixed Effort level ($FE$). The trophic level of $H$, $TL_H$, had a significant but weak effect. Graphs indicate effect across tested ranges and dotted region shows 95% confidence interval. Summary table provides summary statistics. Additional CART results are as follows: folded $R^2 = 0.69$, Highest contributors: Effort=46%, $\log(B_{H0})$ =11%.

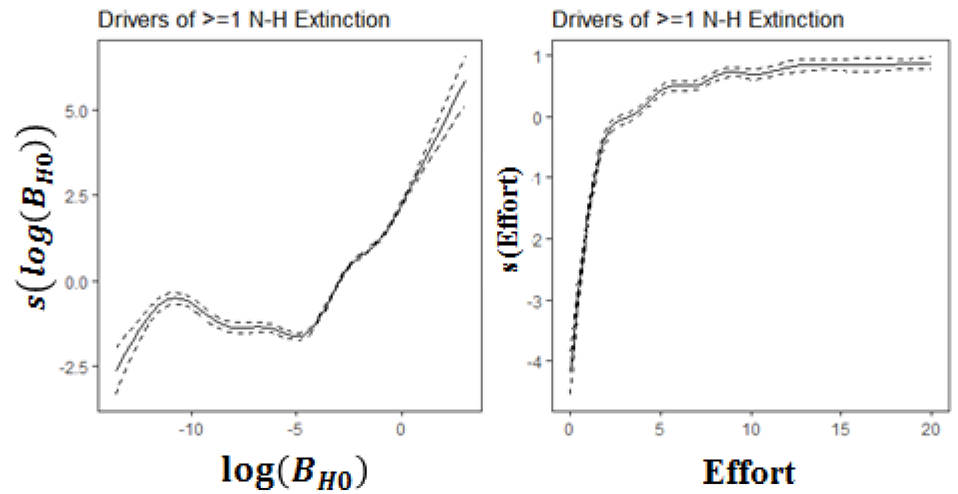


| Effect on Probability of at least 1 Non-Harvested Extinction (Binomial GAM) | | | | | | |
|---|---|---|---|---|---|---|
| Model | p-value | UBRE | Scale Est | $R^2$ | Deviance Explained | AIC |
| $log(B_{H0})$ | $log(B_{H0})$**** | -0.08 | 1 | 0.21 | 18.5% | 35516.1 |
| $FE$ | $FE$**** | 0.06 | 1 | 0.05 | 6.3% | 40828.51 |
| $log(B_{H0}) + FE$ | $log(B_H0)$**** $FE$**** | -0.16 | 1 | 0.29 | 26.3% | 32157.96 |

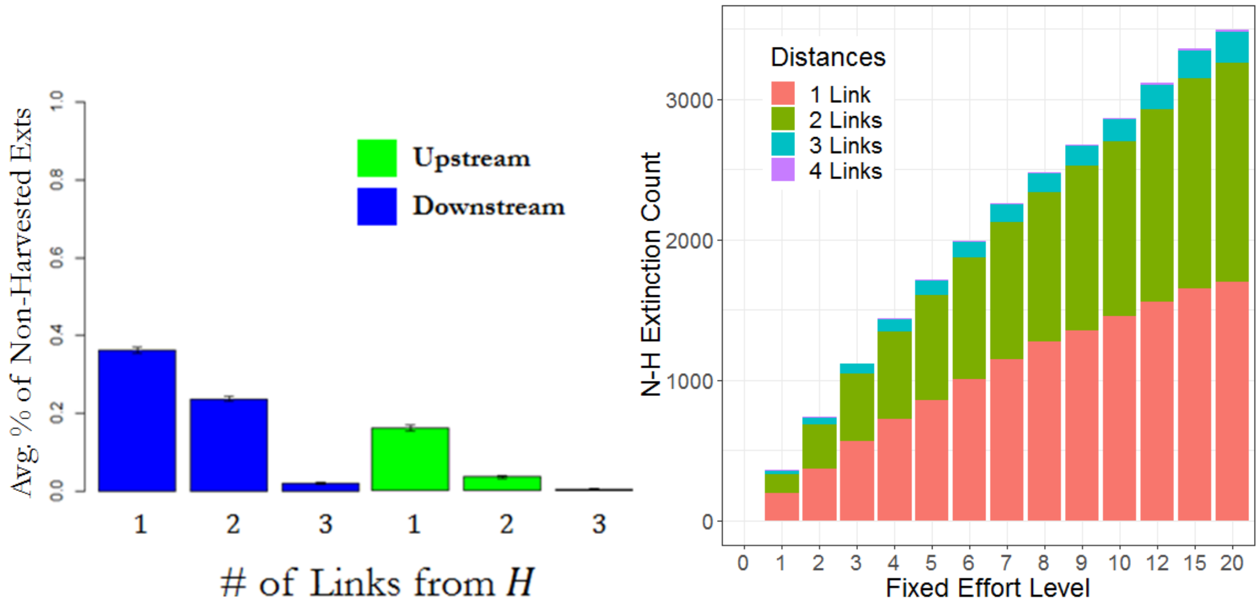


Figure S9: Summary of locations of *N-H* extinctions in food-webs in Fixed Effort simulations. Left: Cumulative count of the trophic distance away from *H* of each *N-H* extinction across all Fixed Effort levels. Trophic distance is defined as the minimum number of trophic links connecting *H* to the extinct *N-H* species. Bars show standard error. Right: The average percent of extinctions up and downstream from *H* and the trophic away from *H* in Fixed Effort simulations.

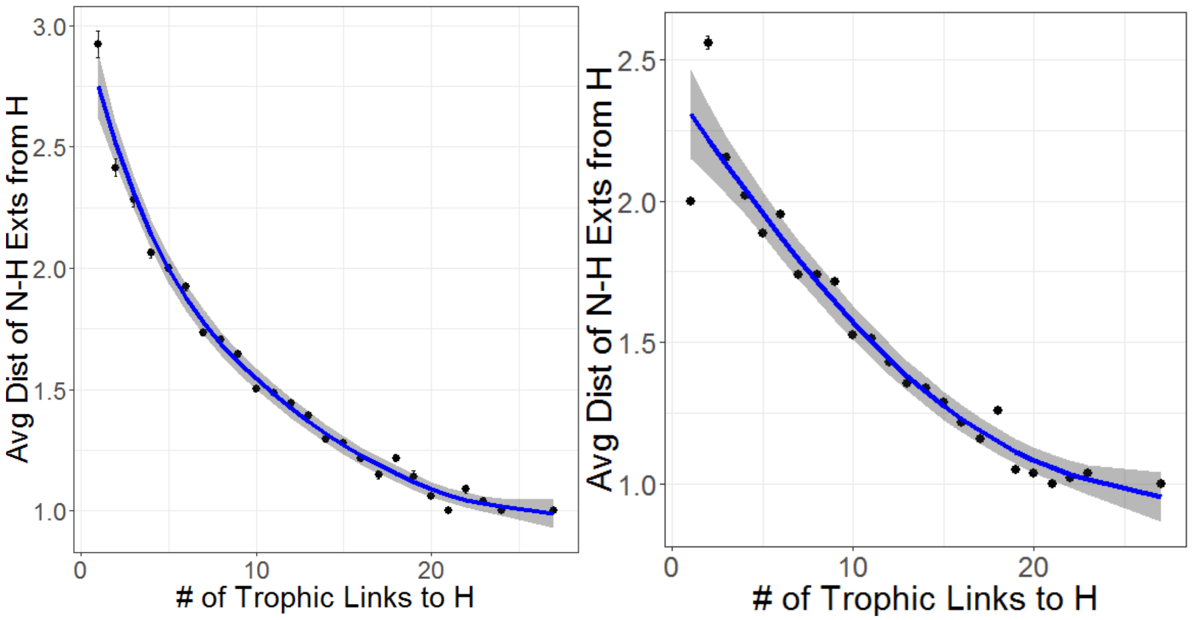


Figure S10: The average distance away from *H* of all *N-H* extinctions from the Fixed Effort simulations (left) and the Open Access simulations (right) as a function of the number of trophic links of *H*. Error bars on points represent standard error. A loess regression (blue) is presented in each graph for visual clarity. Shaded region shows 95% confidence interval.

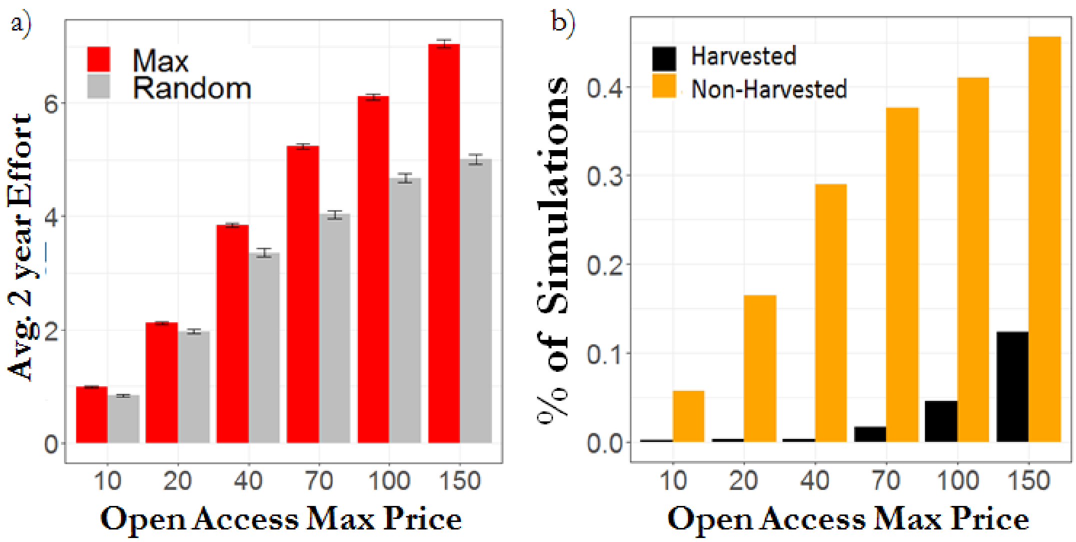


Figure S11: Effects of Open Access Max Price on effort and ecological outcomes. a) Bar graph showing the average Open Access Effort levels across 2 years of fishing at the different tested max price ($a$) values for both the randomly selected fish treatment (gray) and the max abundant fish treatment (choosing H with highest $B_{H0}$; red). Choosing more abundant species to harvest (red) supports higher effort levels. Lines indicate standard error. b) Bar graph showing the percentage of all fishing simulations that resulted in $H$ extinctions (black) and/or $N$-$H$ extinctions (orange) at all tested Open Access Max Price ($a$) values.

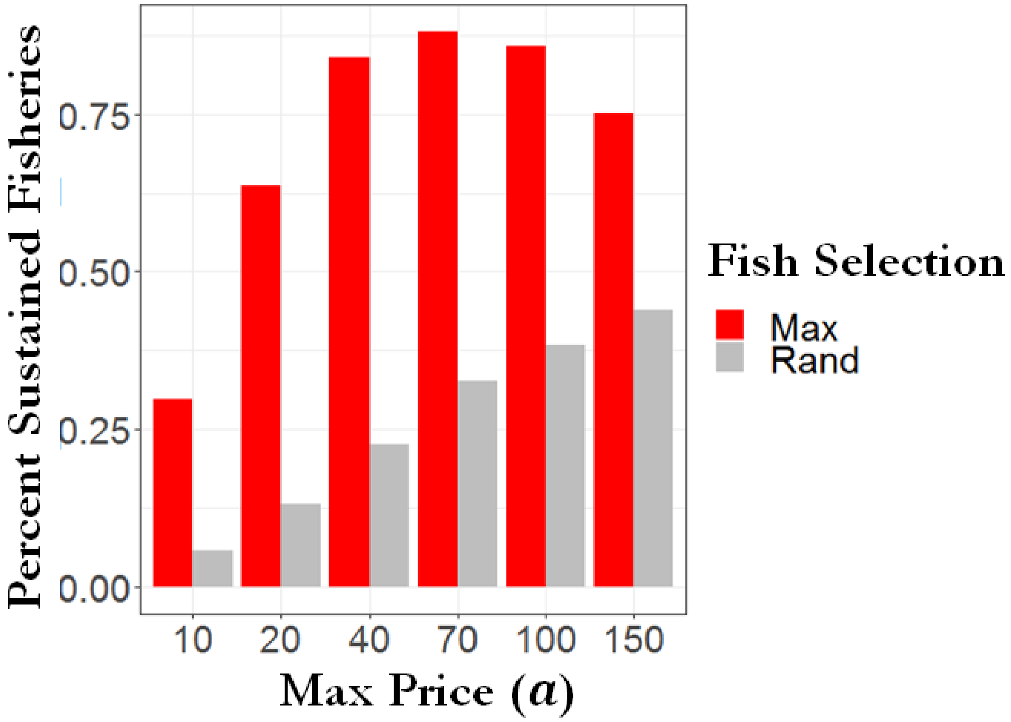


Figure S12: Histogram from Open Access results showing the percent of simulations which produced sustained fisheries across the different max price values (a) and the two fish choice treatments, maximum abundant species (red) and randomly chosen species abundance (grey).

Table S4: Table detailing the statistical support for each driver of Open Access Effort growth through average peak values of Open Access Effort within the first year of fishing. Fits and summary stats are derived from a linear Gaussian model.

| Effect on Average Peak Open Market Effort Value w/in 1st yr (Gaussian linear model) | | | | | | |
|---|---|---|---|---|---|---|
| Model | p-value | Estimate(Std. Er.) | T value | $F_{4\&55829}$ | $R^2$ | AIC |
| $log(B_{H0})$ | $log(B_{H0})$ **** | $log(B_{H0})$:0.89(0.006) | $log(B_{H0})$:147.7 | 2.18e04 | 0.28 | 181994.5 |
| $log(B_{H0}) + \log(a)$ | $log(B_{H0})$**** $\log(a)$**** | $log(B_{H0})$: 1.21(0.004) $\log(a)$:1.21(0.005) | $log(B_{H0})$: 288.2 $\log(a)$: 264.9 | 5.97e04 | 0.68 | <u>136553.4</u> |
| $log(B_{H0}) + \mu$ | $log(B_{H0})$**** $\mu$**** | $log(B_{H0})$:0.91(0.006) $\mu$:1.0(0.014) | $log(B_{H0})$:156.53 $\mu$:68.86 | 1.42e04 | 0.34 | 177445.7 |
| $log(B_{H0}) + b$ | $log(B_{H0})$**** $b$*** | $log(B_{H0})$:0.89(0.006) $b$:-0.09(0.026) | $log(B_{H0})$:147.68 $b$:-3.38 | 1.09e04 | 0.28 | 181985.1 |
| $log(B_{H0}) + \log(a) + \mu + b$ | $log(B_{H0})$**** $\log(a)$**** $\mu$**** $b$*** | $log(B_{H0})$:1.24(0.003) $\log(a)$:1.23(0.004) $\mu$:1.12(0.009) $b$:-.09(0.016) | $log(B_{H0})$:333.11 $\log(a)$:305.85 $\mu$:126.35 $b$:-5.98 | 4.24e04 | 0.75 | **<u>122488.4</u>** |

Table S5: Table detailing the statistical support for each driver of Open Access Effort growth through average peak values of Open Access Effort within the first year of fishing. Fits and summary stats are derived from a Generalized Additive Model (GAM). The starting biomass of $H$ is logged ($\log(B_{H0})$) and treated as a continuous variable while max price ($a$), Effort sensitivity to profit changes ($\mu$), and price sensitivity to yield ($b$) are treated categorically. Each model’s summary statistics and explanatory power is shown in the right table. The fullest model is revealed to be the best model. The effect of each variable in the full model is displayed in the left four panel graphs.

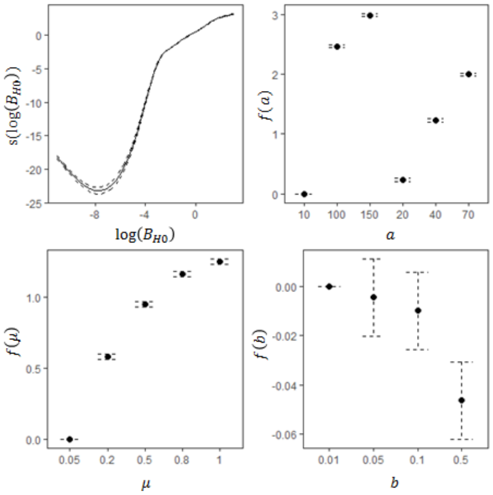


| Effect on Average Peak Open Market Effort Value w/in 1st year (Gaussian GAM) | | | | | | |
|---|---|---|---|---|---|---|
| Model | p-value | GCV | Scale Est | $R^2$ | Deviance Explained | AIC |
| $log(B_{H0})$ | $log(B_{H0})$ **** | 1.44 | 1.44 | 0.32 | 32 | 178877 |
| $log(B_{H0}) + a$ | $log(B_{H0})$**** $a$**** | 0.63 | 0.63 | 0.701 | 70.1 | 132948.2 |
| $log(B_{H0}) + \mu$ | $log(B_{H0})$**** $\mu$**** | 1.31 | 1.31 | 0.378 | 37.8 | 173879.9 |
| $log(B_{H0}) + b$ | $log(B_{H0})$**** $b$*** | 1.44 | 1.44 | 0.32 | 32 | 178866.8 |
| $log(B_{H0}) + a + \mu + b$ | $log(B_{H0})$**** $a$****, $\mu$**** $b$*** | 0.48 | 0.48 | 0.77 | 77.3 | 117528.6 |

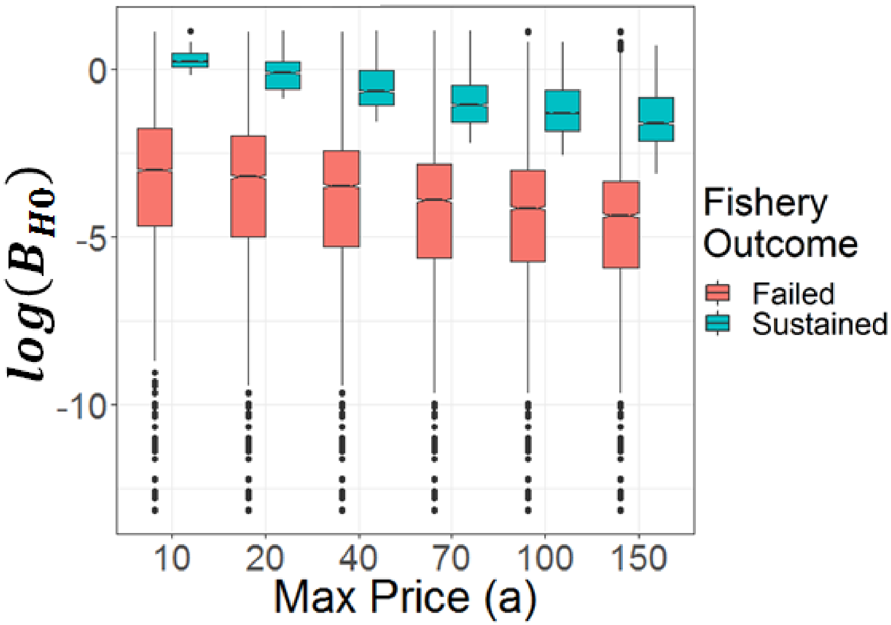


Figure S13: Box and whisker plot detailing the starting biomass of $H$ (i.e. $B_{H0}$) which led to failed fisheries (red) and sustained fisheries (blue) for each tested max price value ($a$) when randomly selecting $H$ in the open market simulations. Boxes represent the interquartile range with the horizontal line showing the median, the lower box representing the 25 percentile, and the upper box showing the 75 percentile. Upper and lower lines extending from the boxes show the most extreme values within 1.5 times the $75^{th}$ and $25^{th}$ percentile respectively. Outliers are shown as single dots.

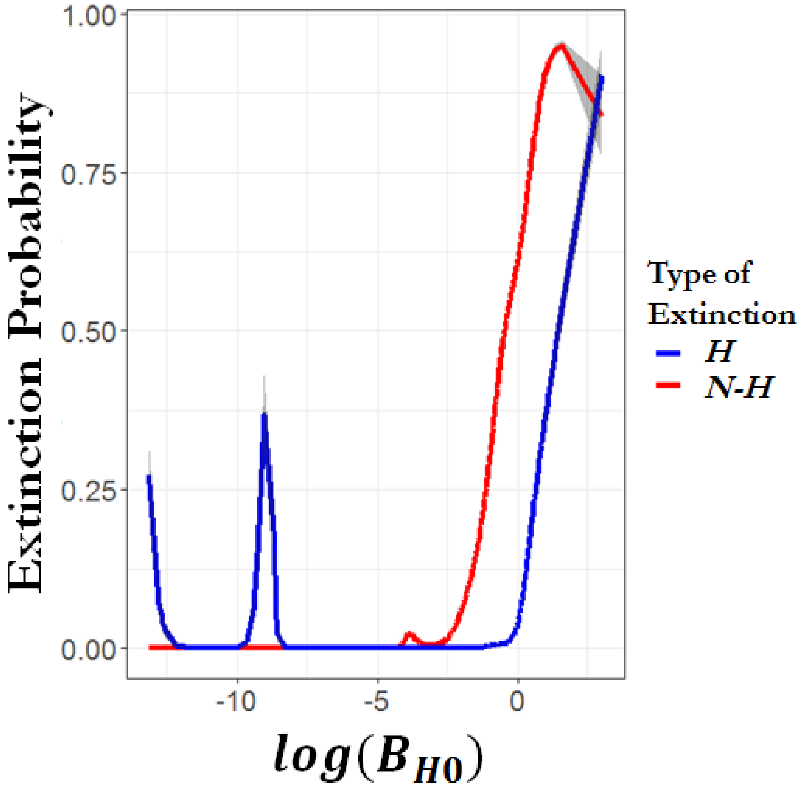


Figure S14: Open Access extinction probabilities across $B_{H0}$. Binomial GAM regressions of the probability of the occurrence of $H$ extinctions (blue) or ≥ 1 $N$-$H$ extinction (red) across the range of starting $H$ population biomass ($B_{H0}$) on a log scale. Shaded region indicates 95% confidence interval. In general, the highest probability of either kind of extinction occurs when harvesting high $B_{H0}$ fish populations. $H$ extinctions do show spikes over limited ranges at smaller levels of $B_{H0}$ despite Open Access fisheries failing at those low levels of $B_{H0}$. At the lowest levels of $B_{H0}$ tested, even the briefest fishing pressure of an initialized fishery is enough to cause H extinctions because $B_{H0}$ is so low. The short spike in $H$ extinctions after $\log(B_{H0}) \approx -\frac{10\mu g}{LC}$ occurs because those levels of $B_{H0}$ are enough to briefly support fishing effort before failing. This limited fishing effort does cause some $H$ extinctions in some webs. Higher levels of $B_{H0}$ allows for $H$ to withstand the fishing pressure before fishery collapse, which is why the probability of $H$ extinctions does eventually drop to zero again.

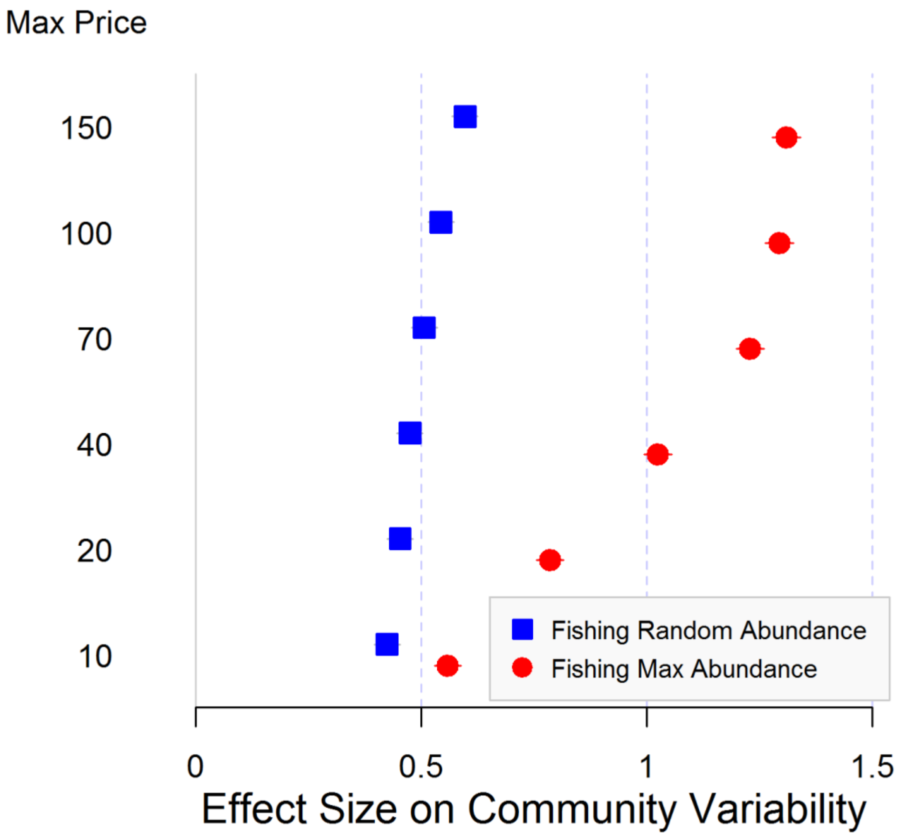


Figure S15: Hedge’s G comparison of the mean community stability of food webs fished through the Open Access management and the closest Fixed Effort value across the six max price values tested. Mean community stability is the average C.V. of each population’s biomass after fishing starts. Dots represent effect size and lines show 95% confidence intervals. The 0 value indicates no difference between Fixed Effort and Open Access treatments and higher magnitude values indicate more variability in the Open Access treatments. Results indicate that both Open Access fish choice treatments create more variation in community biomass with a greater effect seen when fishing the maximum abundant fish.

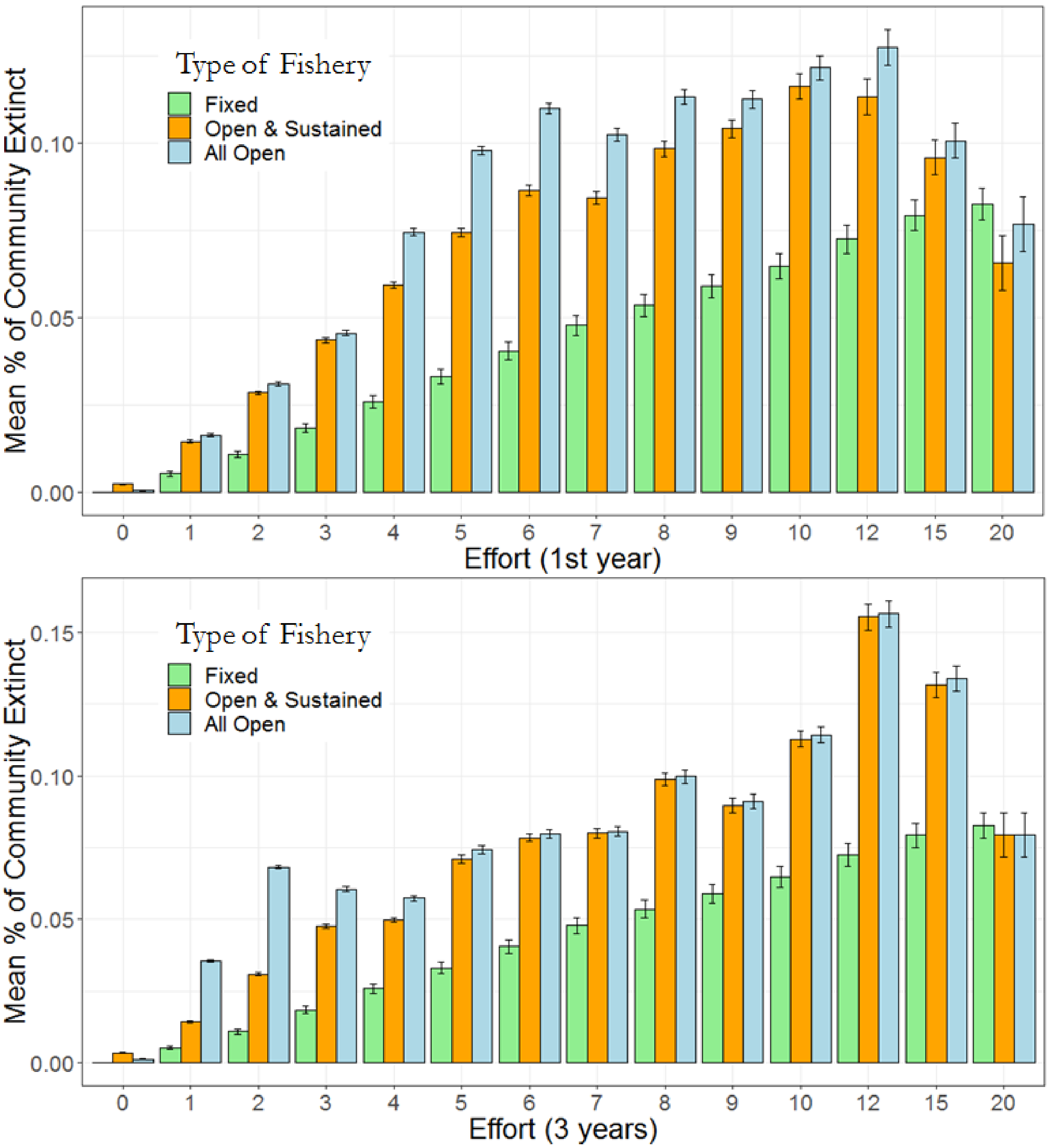

Figure S16: Mean percentages of species lost to *N-H* extinctions when Open Access treatments give comparable effort levels to tested Fixed Effort values. Open Access levels are taken as averages of effort across (top) 1 year and (bottom) 3 years. *N-H* extinction percentages are given for all Open Access simulations (light blue) and only the Open Access simulations that produced a sustained fishing effort (orange). Fixed Effort results are shown in light green). Bars represent standard error.

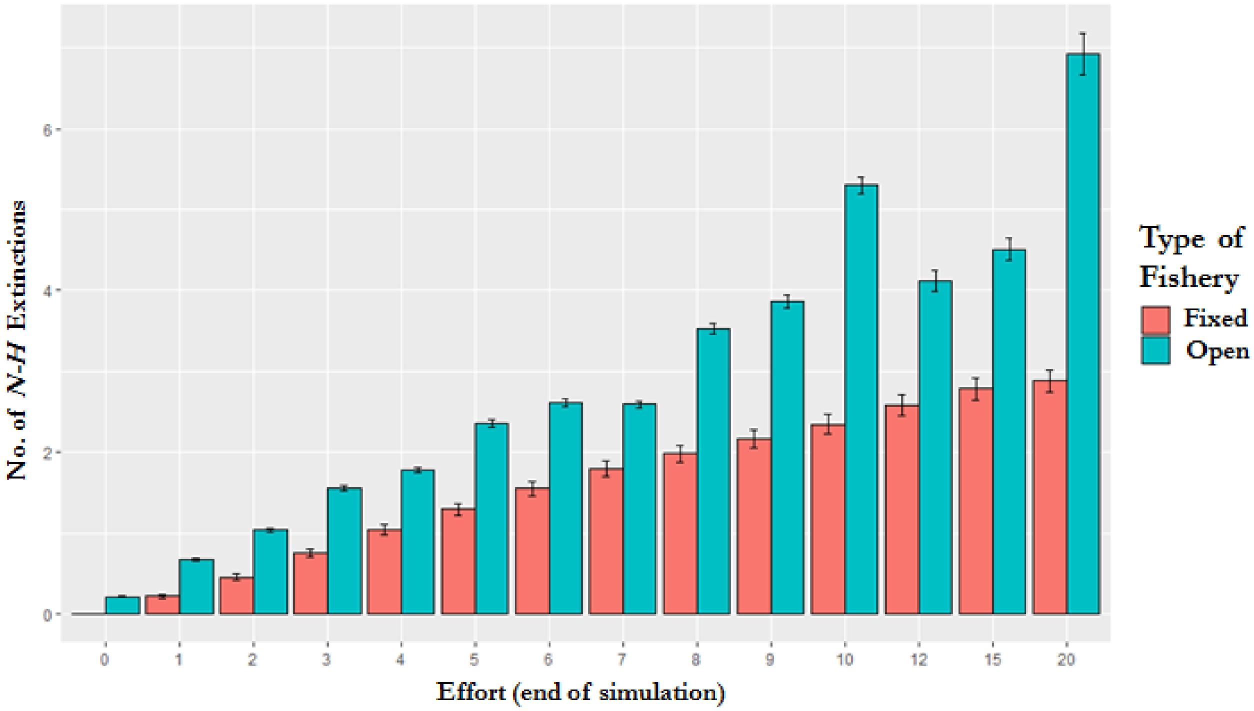


Figure S17: Bar graph comparing the number of *N-H* extinctions in Fixed Effort fisheries to Open Access fisheries with comparable effort levels at t=8000. Bars represent standard error.

Permissions for reproduction of Figure 4 from Smith et al 2011 have not yet been acquired. Upon obtaining permissions, figure will be reproduced here for ease of readers.

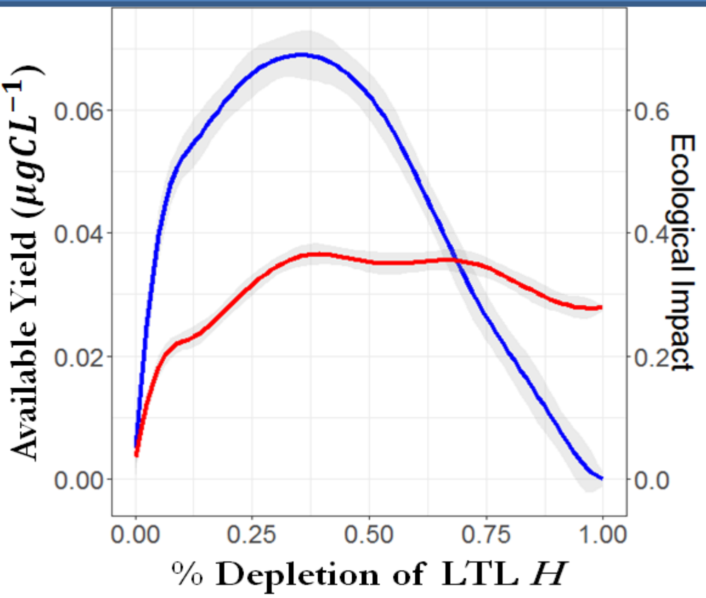


Figure S18:

***__**Top:**__** A recreation of Figure 4 from Anthony Smith et al 2011 (23), Impacts of Fishing Low–Trophic Level Species on Marine Ecosystems. The paper investigates, among other things, the ecosystem impact of harvesting/fishing low trophic level (LTL) species. Smith et al details how the level of LTL species depletion due to fishing affects potential yield (as a function of Maximum Sustainable Yield (MSY)). It also shows the ecological impact of each level of LTL species depletion through a measure of the proportion of other species groups' which have biomass effected by ≥ 40%.

__**Bottom:**__ Using the statistical power from the FE treatment which harvested every possible fish, our model framework was used to qualitatively reproduce the above figure from Smith et al 2011. The exact trophic level cut off for being considered a LTL species is not listed in Smith et al 2011, but the species included in Smith et al's analysis are described as plankton feeders so we considered LTL species from our results as H species with a trophic level ≤ 3. Though, results do not qualitatively change with slight increases in the LTL cutoff value. This cutoff gave 737 fish populations across 65.6% of our webs. The x-axis of LTL depletion was created using the % of $H$ biomass directly depleted by Fixed Effort fishing. While we do not calculate the MSY for each harvested species we can calculate the resulting potential yield at each level of depletion by multiplying the remaining biomass of $H$ by the Fixed Effort level and the catchability coefficient ($q$). This gives the same negative parabola shape (shown in blue: GAM result with shaded region showing 95% confidence interval) and would do the same as a percentage of MSY. Ecological impact on the right axis is calculated in the same manner used in Smith et al 2011 (red; GAM result with shaded regions showing standard error). While the ecological impact shows a qualitatively similar increase, it rises fasters and seems to hit an asymptotic value. This is due to our food webs being capped at 30 species per web which makes it easier to affect a higher percent of species with less absolute numbers.

*****Please note: we plan on contacting AAAS regarding permission to reuse Figure 4 from Smith et al 2011. We will purchase the rights to reuse the figure given Reviewer acceptance of its place in this manuscript’s SI and the general acceptance of the manuscript.**
